# Aryl hydrocarbon receptor (AhR) restrains the maintenance and acquisition of the anti-inflammatory identity of monocyte-derived macrophages (M-MØ)

**DOI:** 10.64898/2026.11.11.765994

**Authors:** Alejandro del Castillo Buey, Concha Nieto, Israel Ríos, Miriam Simón-Fuentes, Cristina Herrero, Ángel Maldonado, Alba Angelina, Ángela Sanzo-Machuca, Oscar Palomares, Azucena Salas, Miguel A. Vega, Ángel L. Corbí

**Affiliations:** Myeloid Cell Laboratory, Centro de Investigaciones Biológicas, CSIC, Madrid, Spain; Department of Biochemistry and Molecular Biology, School of Chemistry, Complutense University of Madrid, Madrid, Spain; Inflammatory Bowel Disease Unit, Institut d’Investigacions Biomèdiques August Pi I Sunyer (IDIBAPS), Hospital Clínic, Barcelona, Spain; Centro de Investigación Biomédica en Red de Enfermedades Hepáticas y Digestivas (CIBEREHD), Spain; Universitat de Barcelona (UB), Barcelona, Spain

**Author notes:** **Corresponding author**: Dr. Ángel L. Corbí. Centro de Investigaciones Biológicas “Margarita Salas”, CSIC. Ramiro de Maeztu, 9. Madrid 28040.

**Keywords:** Macrophages, Inflammation, Transcription Factors

## Abstract

The Aryl hydrocarbon receptor (AhR) is an essential environmental sensor bridging metabolic cues with immune regulation, yet its precise role in defining human monocyte-derived macrophage identity remains unclear. Here, we demonstrate that AhR functions as a critical transcriptional rheostat that selectively restrains the anti-inflammatory and pro-fibrotic identity of M-CSF-dependent monocyte-derived macrophages (M-MØ). Integrated ChIP-Seq and transcriptomic profiling revealed that AhR activity is enriched in M-MØ in a MAFB-dependent manner, directly occupying regulatory regions of core M-MØ identity genes, including *CD163*, *LGMN*, and *IL10*. Unexpectedly, siRNA-mediated knockdown and CRISPR/Cas9 ablation demonstrated that AhR primarily functions as a repressor; its deficiency accelerated and exacerbated the acquisition of a “resident-like” YVE1^+^/FOLR2^+^) and pro-fibrotic phenotype. Functionally, AhR restricted the M-MØ anti-inflammatory response to TLR7 ligation, and limited an overly tolerogenic profile during M-MØ-driven T-cell responses. Pharmacological modulation using the endogenous ligand FICZ and the antagonist SR1 confirmed that AhR activity limits the magnitude of the MAFB-driven program and the autocrine IL-10 signaling axis. Collectively, these findings establish AhR as a fundamental checkpoint that prevents macrophages from acquiring an excessive anti-inflammatory state, thereby ensuring balanced responses essential for tissue homeostasis and immune surveillance.

## INTRODUCTION

Macrophages exhibit a substantial functional heterogeneity dictated both by their origin and their microenvironmental context ^1^. Tissue-resident macrophages (TRM) perform core homeostatic functions (tissue remodeling, repair and maintenance) in response to the prevailing extracellular cues ^1^ while executing organ-specific roles across distinct TRM subsets ^2^ ^3^ ^4^ ^5^ ^6^ ^7^ ^8^ ^9^. Although most TRM originate during embryonic development, TRM in certain tissues (*e.g*., gut) are mostly replenished by circulating monocytes, with the relative proportions of embryonically-derived TRM and homeostatic monocyte-derived macrophages (MDM) remaining tissue-specific ^1^. During inflammation, disease-associated signals recruit monocytes that differentiate non-homeostatically into MDM ^110^. These inflammatory MDMs, generated under the influence of tissue injury cues and pro-inflammatory cytokines ^11^ ^12^ ^13^, are functionally and phenotypically distinct from monocyte-derived TRM ^1^, and facilitate pathogen clearance, inflammation resolution, tissue repair and return to homeostasis ^1^. However, depletion of homeostatic TRM alongside pathological MDM infiltration drives disease progression ^1^ in severe COVID-19 ^14^ ^15^, inflammatory bowel disease, and cardiovascular, neurodegenerative, hepatic, and neoplastic conditions ^16^ ^17^ ^18^ [reviewed in ^1^].

Constitutive homeostatic MDM differentiation follows distinct kinetics compared to inflammation-driven differentiation, which induces immediate transcriptional shifts in recruited monocytes ^13^. The functional differences between homeostatic MDM and inflammatory MDM are mirrored *in vitro* by differentiating monocytes with M-CSF (yielding homeostatic, anti-inflammatory, immunosuppressive M-MØ) or GM-CSF (resulting in pro-inflammatory GM-MØ) ^19^ ^20^. Indeed, M-MØ and GM-MØ exhibit very distinct transcriptomic programs ^19^ ^20^: M-MØ resemble *in vivo* homeostatic MDM, pro-fibrotic MDM ^14^ ^15^ ^21^ and pro-tumoral “large tumor-associated macrophages (TAM)” ^22^, whereas GM-MØ resemble inflammatory MDM ^14^, anti-tumoral “small TAM” ^22^ and lung alveolar macrophages ^23^. In M-MØ, this core anti-inflammatory transcriptional profile depends primarily on the MAFB transcription factor ^20^ ^21^.

The Aryl hydrocarbon receptor (AhR), a basic helix-loop-helix PER-ARNT-SIM transcription factor ^24^, senses exogenous xenobiotics, dietary ligands, and tryptophan-derived bacterial metabolites ^24^ ^25^. Although AhR is broadly expressed in myeloid cells, its role in macrophage specialization and activation remains controversial due to ligand-dependent and context-specific effects [reviewed in ^24^]. AhR activation has been reported to enhance transcription in GM-MØ but not in M-MØ ^26^, and AhR signaling in monocytes or murine bone-marrow-derived precursors can direct differentiation toward dendritic cells rather than macrophages ^27^ ^28^. Conversely, we previously demonstrated that serotonin upregulates the expression of M-MØ-enriched genes in an AhR-dependent manner ^29^. Regarding macrophage activation, *in vitro* and *in vivo* studies indicate that deletion or inhibition of AhR favors the acquisition of a proinflammatory phenotype ^24^, and that AhR prolongs the duration of the IL-4-induced STAT6 activation in monocytes through non-genomic mechanisms ^30^.

To clarify the precise role of AhR in human MDM specialization and activation, we evaluated genetic and pharmacological AhR inhibition in M-CSF-dependent monocyte-derived macrophages (M-MØ). Here, we report that AhR activity and target gene expression in macrophages depends on the inflammatory state, that AhR transcriptional activity is enriched in anti-inflammatory M-MØ, and that AhR restrains both the acquisition and the maintenance of the anti-inflammatory phenotype of M-MØ.

## RESULTS

### 1.- AhR activity and target gene expression are selectively enriched in anti-inflammatory macrophages in a MAFB-dependent manner

To determine the contribution of AhR to the transcriptional and functional landscape of human macrophages, we first evaluated its expression and basal activity across monocytes, anti-inflammatory M-MØ, and pro-inflammatory GM-MØ (Figure 1A). Although *AHR* mRNA was highest in uncommitted monocytes (Supplementary Figure 1A) and subcellular localization was comparable between macrophage subsets (Supplementary Figure 1B), M-MØ displayed significantly higher AhR protein levels and basal transcriptional activity than their GM-MØ counterparts (Figure 1B-C). This activity was further enhanced by the canonical agonist FICZ ^31^ and effectively blocked by the antagonist CH223191 (Figure 1C). Transcriptomic analysis revealed that the M-MØ profile was strongly enriched for AhR-activated gene signatures identified across independent datasets generated through pharmacological activation [Benzo[a]pyrene, BaP, ^32^; VAF347 ^33^; indole-3-carbinol, I3C, ^34^], antagonism [StemRegenin1, SR1, ^35^], or genome-wide ChIP-Seq mapping ^36^ (Figure 1D-E; Supplementary Figure 1C-D) ^19^. Notably, the expression of these AhR-dependent gene sets in M-MØ strictly required MAFB, the master transcription factor governing M-MØ identity ^20^ ^21^ ^37^ (Figure 1D). Conversely, GM-MØ were enriched for genes typically repressed by AhR ^32^ ^33^ (Figure 1D), a pattern validated in independent M-MØ and GM-MØ datasets ^38^ ^39^ (Figure 1F; Supplementary Figure 1E). Furthermore, AhR antagonism (CH-223191, SR1) impaired the expression of M-MØ-enriched hallmark AhR targets (Figure 1G) whose proximal regulatory regions were directly occupied by AhR, as confirmed by genome-wide ChIP-Seq mapping in M-MØ (Figure 1H; Supplementary Figure 1F-G). Highlighting the *in vivo* relevance of these results, AhR-regulated genes were preferentially expressed in MAFB-dependent homeostatic gut macrophage subsets (M2, M2.2) ^9^ (Figure 1I). Collectively, these data demonstrate that AhR transcriptional activity, and its associated transcriptome, is selectively enriched in M-CSF-dependent M-MØ, where MAFB contributes to the preferential expression of canonical AhR target genes.

**Figure 1.**
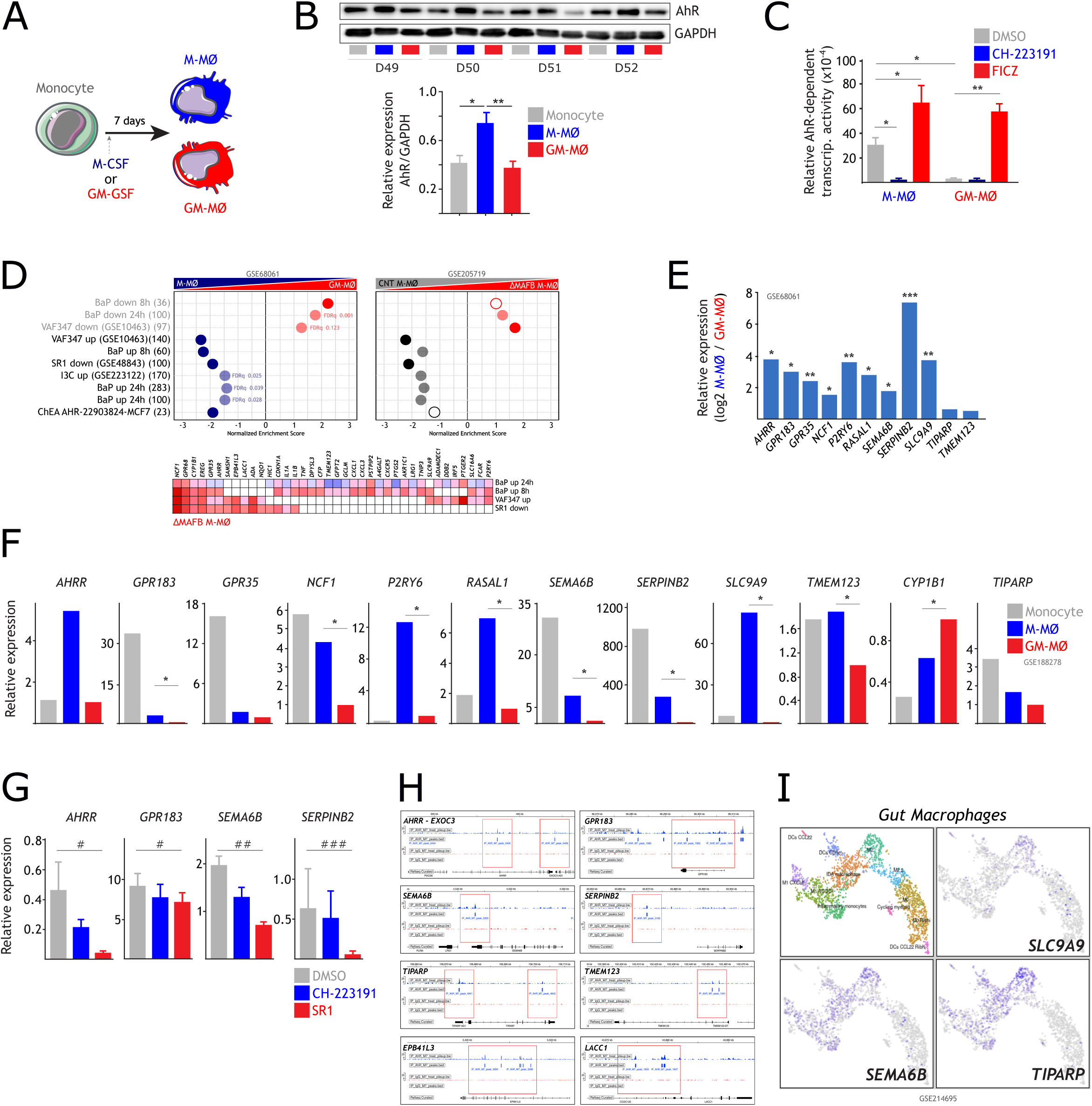
– AhR expression and activity are enriched in the anti-inflammatory M-MØ lineage. **A**. Schematic workflow for generation of monocyte-derived anti-inflammatory (M-MØ) and pro-inflammatory (GM-MØ) macrophages. **B**. Western blot of AhR protein in monocytes, M-MØ, and GM-MØ (n=4) (*Upper panel*). GAPDH served as protein loading control. Bar graph shows AhR/GAPDH protein ratios (mean ± SEM; *, p<0.05; **, p<0.01) (*Lower panel*). **C**. AhR transcriptional activity in M-MØ and GM-MØ treated with DMSO, FICZ or CH-223191, determined by Xenobiotic Response Element (XRE)-reporter luciferase assay relative to a constitutive CMV promoter driving *Renilla* luciferase expression (n=4; mean ± SEM; *, p<0.05; **, p<0.01). **D**. GSEA of canonical AhR-regulated gene sets [from ^3233343536^] on the ranked comparison of GM-MØ vs. M-MØ transcriptome (GSE68061) (*Left panel*) or Control M-MØ (CNT M-MØ) vs. ΔMAFB-M-MØ transcriptome (GSE205719) (*Right panel*). Leading-edge analysis of AhR-regulated genes affected by *MAFB* is shown (*Lower panel*). Normalized Enrichment Score (NES) and FDRq values are indicated (FDRq=0, solid colors; FDR<0.25, light colors; Not significant, empty circles). **E**. Relative expression of AhR-regulated genes in GM-MØ and M-MØ (GSE68061) (*, p<0.05; **, p<0.01; ***, p<0.005). **F**. RNA-Seq validation of relative mRNA levels of the indicated genes in monocytes, M-MØ and GM-MØ (GSE188278) (n=3; *, adjp<0.05). **G**. qRT-PCR analysis of AhR-target genes in M-MØ treated with DMSO, CH-223191 or StemRegenin1 (SR1) (n=3 for *GPR183, SEMA6B*; n=4 for *AHRR, SERPINB2;* mean ± SEM; ^#^, p<0.05; ^##^, p<0.01). **H**. ChIP-seq profiles showing AhR occupancy at proximal regulatory regions of the indicated genes in M-MØ. **I**. Expression of AhR-regulated genes in homeostatic gut macrophage subsets *in vivo* (M2, M2.2) (GSE214695) ^9^.

### 2.- AhR restricts the full anti-inflammatory gene and cytokine profile of M-MØ

Gene ontology analysis of 1008 genes exhibiting AhR binding at proximal regulatory regions (−1000/+100) demonstrated enrichment of genes co-expressed with AhR and overexpressed in specific macrophage subsets (Supplementary Figure 1H). AhR directly bound proximal regulatory regions of core M-MØ markers, including *ADGRE1*, *CCL2*, *CCL8*, *CD163*, *CD163L1*, *IL10*, *LGMN*, *MAFB*, *SIGLEC1*, *SLC40A1,* and *STAB1* ^19^ ^20^ (Supplementary Figure 1I). To determine functional consequences, we generated AhR-deficient macrophages using siRNA (ΔAHR M-MØ) (Figure 2A). AhR knockdown caused extensive transcriptional remodeling (Figure 2B), with reduced expression of canonical AhR-dependent genes ^35^ (Supplementary Figure 2A-C) and, unexpectedly, upregulated genes with the highest proximal AhR binding (302-geneset, Supplementary Figure 2D-E), indicating a predominantly repressive role for AhR. Regarding the global M-MØ transcriptional signature ^19^ ^20^, ΔAHR M-MØ exhibited an enhanced M-MØ-specific anti-inflammatory program and a diminished GM-MØ-associated pro-inflammatory signature (Figure 2C). ΔAHR M-MØ showed significant enrichment for core M-MØ identity markers (*Odds ratio, 8.06; p-value, 1.03×10^-^*^15^), including *CCL2*, *CCL8*, *CD163*, *LGMN*, and *SLC40A1* (Figure 2D), all of which exhibit direct AhR binding to their proximal regulatory regions (Supplementary Figure 1I). This inhibitory role was validated at the protein level, as AhR deficiency increased CD163 expression and elevated secretion of CCL8, LGMN, IL-10, and CXCL10 (Figure 2E-F), thus establishing AhR as a limiting factor for the expression of the genes that define the M-MØ identity.

**Figure 2.**
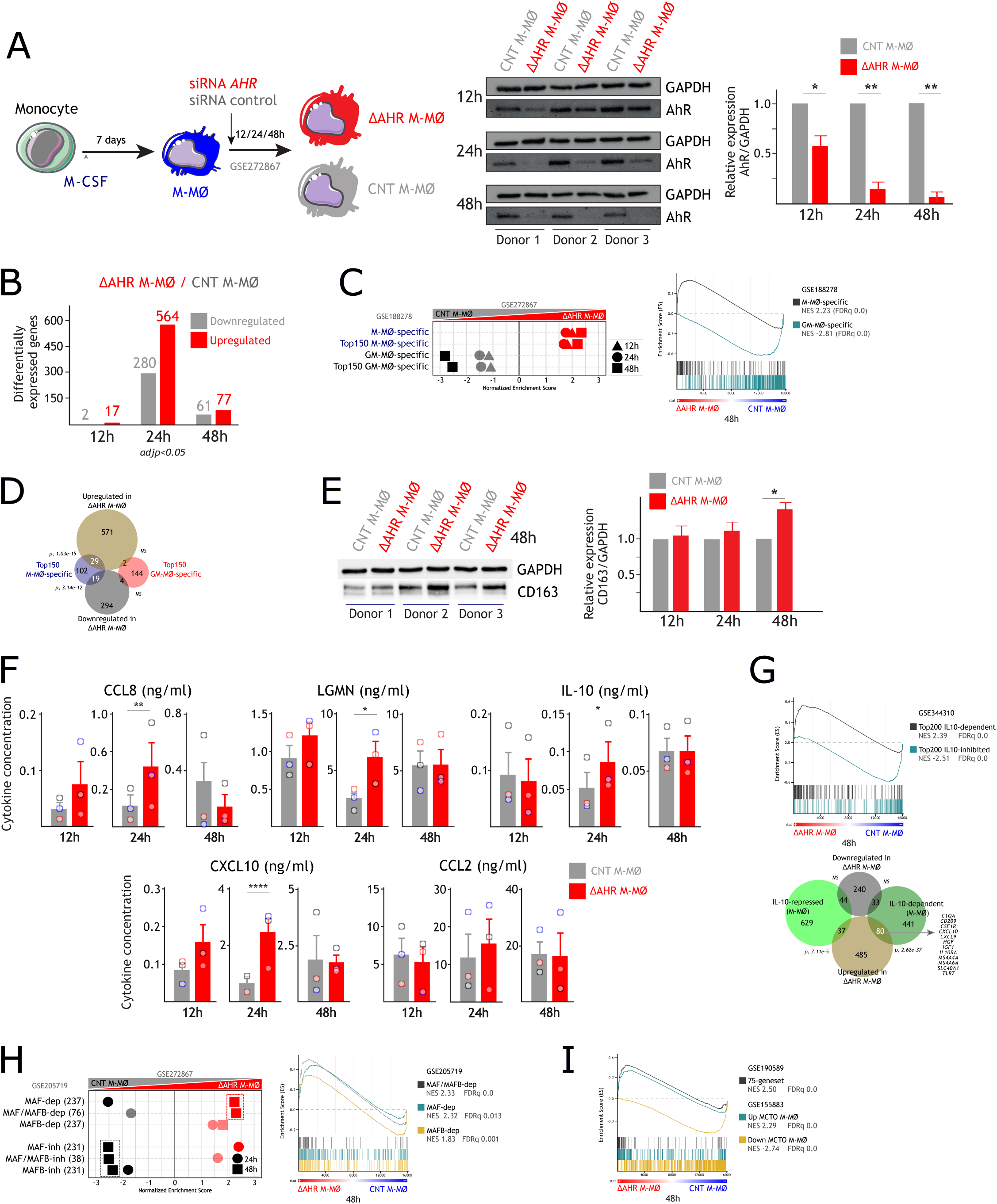
– AhR restrains the core anti-inflammatory profile of monocyte-derived M-MØ. **A**. Schematic representation of the procedure for siRNA-mediated AHR knock-down in monocyte-derived anti-inflammatory (M-MØ) macrophages (*Left panel*). Time-course analysis of AhR expression in M-MØ transfected with a control siRNA (CNT M-MØ) or AHR-specific siRNA (ΔAHR-M-MØ), determined by Western blot (*Middle panel*) and protein quantification (AhR/GAPDH protein ratios, *Right panel*) (n=3; mean ± SEM; *, p<0.05; **, p<0.01). **B**. Quantification of differentially expressed genes between ΔAHR-M-MØ and CNT M-MØ at 12h, 24h and 48h after transfection. **C**. GSEA summary (*Left panel*) and representative profiles (*Right panel*) of M-MØ– or GM-MØ-specific gene sets ^67^ on the ranked comparison of CNT M-MØ vs. ΔAHR-M-MØ transcriptomes at 12, 24 and 48h after silencing. NES and FDRq are indicated (FDRq=0, solid colors; FDR<0.25, light colors). **D**. Overlap of ΔAHR-M-MØ modulated genes (adjp<0.05 at any time point) and top M-MØ– or GM-MØ-specific markers. **E**. Western blot (*Left panel*) and quantification (CD163/GAPDH protein ratios, *Right panel*) of CD163 protein levels in CNT M-MØ and ΔAHR-M-MØ at the indicated time points after transfection (n=3; mean ± SEM; *, p<0.05). **F**. ELISA quantification of the indicated factors released from CNT M-MØ and ΔAHR-M-MØ at the indicated time points after transfection (n=3; mean ± SEM; *, p<0.05; **, p<0.01; ****, p<0.001). **G**. GSEA profiles of IL-10-dependent and IL-10-repressed genes in M-MØ (GSE344310) on the ranked comparison of the CNT M-MØ vs. ΔAHR-M-MØ transcriptome at 48h after silencing (*Upper panel*). NES and FDRq are indicated in each case. Overlap of ΔAHR-M-MØ modulated genes (adjp<0.05 at any time point) and IL-10-dependent or IL-10-repressed genes in M-MØ (*Lower panel*). Significance of each overlap is indicated. **H**. Summary of GSEA, and representative profiles, of MAFB– and MAF-dependent regulons ^41^ on the ranked comparison of the CNT M-MØ vs. ΔAHR-M-MØ gene profiles at the indicated times after silencing. NES and FDRq are indicated (FDRq=0, solid colors; FDR<0.25, light colors). **I**. GSEA profiles of genes up/downregulated in MCTO M-MØ ^68^, or *bona fide* MAFB-dependent genes (75-geneset) ^21^, on the ranked comparison of the CNT M-MØ vs. the ΔAHR-M-MØ transcriptome 48h after silencing. NES and FDRq are indicated in each case.

The augmented production of IL-10 by ΔAHR M-MØ is of particular mechanistic importance because AhR deficiency broadly upregulated IL-10-induced M-MØ genes while downregulating IL-10-repressed genes (GSE344310) (Figure 2G). These upregulated targets included functionally relevant hallmark M-MØ markers like *C1QA*, *CSF1R*, *CXCL10*, *IGF1* and *TLR7* ^20^ ^21^ ^37^ (Figure 2G), implying that AhR contributes to the maintenance of the M-MØ gene signature by limiting the magnitude of its anti-inflammatory program. Since the M-MØ transcriptome is primarily governed by MAFB and MAF ^20^ ^21^ ^37^, we also analyzed the effect of AhR deficiency on these specific regulons ^40^. The ΔAHR M-MØ transcriptome was enriched for MAFB– and MAF-dependent regulons ^41^ ^21^ (Figure 2H; Supplementary Figure 2F), and showed a high overlap with the transcriptome of M-MØ from Multicentric Carpo-tarsal Osteolysis (MCTO) patients, who exhibit augmented MAFB protein levels ^42^ ^43^, and the MAFB ChIP-Seq cistrome ^21^ (75-geneset) (Figure 2I). Collectively, these data indicate that AhR restricts the M-MØ profile by impairing IL-10 expression and the magnitude of the MAFB-driven program. Since MAFB and MAF protein levels were unchanged in AHRM-MØ (Supplementary Figure 2G), the AhR inhibitory effect might take place via functional interference or co-repressor recruitment rather than through direct transcriptional suppression of MAF/MAFB.

### 3.- AhR restrains the *in vivo* signatures of pro-fibrotic, anti-inflammatory, and pro-tumoral macrophages

The M-MØ gene profile closely mirrors several specialized *in vivo* macrophage populations, including homeostatic M2-like gut macrophages (M2, M2.2) ^9^, pathogenic monocyte-derived lung macrophages in severe COVID-19 (MoAM3, CD163^+^/LGMN^+^) ^44^ ^15^, and pro-tumoral “Large TAM” in colorectal liver metastasis ^39^ ^21^ ^22^ (Supplementary Figure 2H-I). Gene Set Enrichment Analysis (GSEA) showed that AhR knockdown significantly enhanced the expression of genes defining homeostatic gut M2/M2.2 subsets, pro-fibrotic MoAM3 and CD163^+^/LGMN^+^ lung macrophages ^44^ ^15^, and Large TAM from both colorectal liver metastasis ^22^ and breast cancer ^45^ (Figure 3A-B; Supplementary Figure 2I-K). Gene ontology analysis of genes shared between AHRM-MØ and these *in vivo* subsets showed strong enrichment for IL-10-dependent networks (Figure 3A-B; Supplementary Figure 2J). Conversely, ΔAHR M-MØ displayed reduced expression of genes characteristic of pro-inflammatory M1-like gut macrophages ^9^, GM-CSF-dependent lung-resident alveolar macrophages (TRAM1, TRAM2, AMØ1) ^4415^, and anti-tumoral “Small TAM” ^39^ ^21^ ^22^ (Figure 3A-B; Supplementary Figure 2H-I). Collectively, these results indicate that AhR activity selectively restrains the *in vivo* signatures of anti-inflammatory, pro-tumoral and pro-fibrotic macrophage phenotypes.

**Figure 3.**
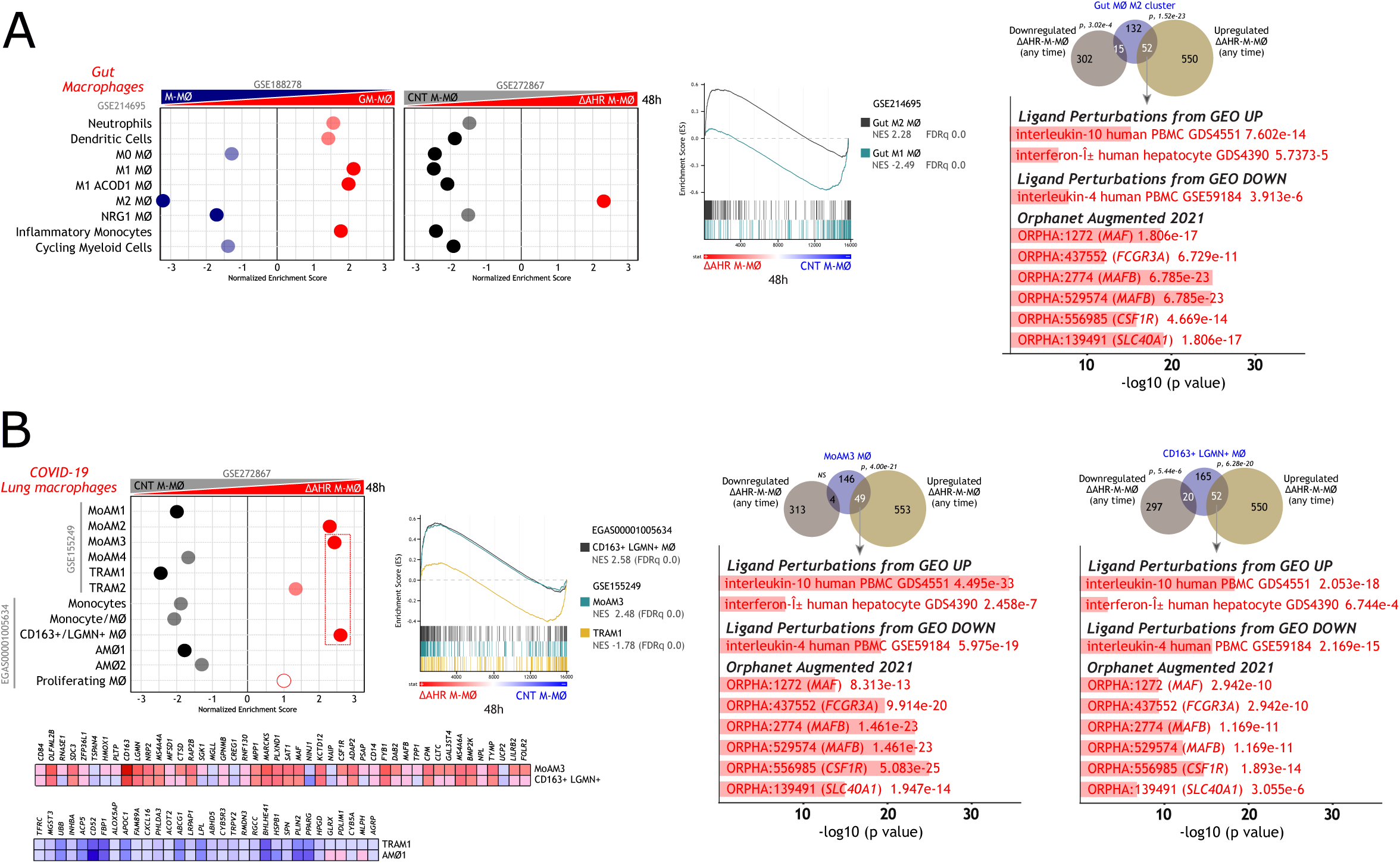
AhR deficiency enhances homeostatic and pathogenic macrophage signatures *in vivo*. **A**. GSEA summary and representative profiles (*Left/center panels*) of the gene sets defining macrophage subsets in human gut ^9^ on the ranked comparison of the CNT M-MØ vs. the ΔAHR-M-MØ transcriptome 48h after silencing. NES and FDRq are indicated (FDRq=0, solid colors; FDR<0.25; Not significant, empty circles). Overlap of ΔAHR-M-MØ modulated genes (adjp<0.05 at any time point) and genes within the Gut M2 MØ cluster is shown (*Left panel*). Gene ontology analysis (Enrichr) ^61^ of the most significant overlap is shown. **B**. GSEA summary, representative profiles and leading-edge analysis (*Left panels*) of the gene sets that define macrophage subsets in severe COVID-19 ^4415^ on the ranked comparison of the CNT M-MØ vs. the ΔAHR-M-MØ transcriptome 48h after silencing. NES and FDRq are indicated (FDRq=0, solid colors; FDR<0.25; Not significant, empty circles). Overlaps of ΔAHR-M-MØ modulated genes (adjp<0.05 at any time point) and genes within the MoAM3 MØ cluster or the CD163+LGMN+MØ cluster are shown (*Right panel*). Gene ontology analysis (Enrichr) ^61^ of the most significant overlaps are shown.

### 4.- Pharmacological modulation confirms AhR-mediated repression of the M-MØ transcriptome

Given the enrichment of the M-MØ profile upon AhR silencing, we assumed that ligand-induced AhR activation would yield the reciprocal outcome. To test this, M-MØ were treated with the endogenous agonist FICZ (FICZ-M-MØ) or the antagonist SR1 (SR1-M-MØ) for 24 hours (Figure 4A-B). FICZ induced proteasomal degradation of AhR (Supplementary Figure 3A), a hallmark of sustained AhR activation ^46^ ^47^ ^48^ ^49^, upregulated canonical AhR-target genes characterized ^32^ ^33^ ^34^ ^35^ ^36^ (Figure 4B; Supplementary Figure 3B) and, more importantly, downregulated core M-MØ markers (*CD163, FOLR2, LGMN, SLC40A1*) alongside MAFB-dependent genes (Figure 4C-D; Supplementary Figure 3C-D). Cross-referencing ΔAHR M-MØ and FICZ-M-MØ transcriptomes defined a core set of 109 “AhR-dependent” genes (downregulated by silencing, upregulated by FICZ) and 169 “AhR-repressed” genes (upregulated by silencing, inhibited by FICZ) (Figure 4E; Supplementary Figure 3E). SR1 treatment partially mirrored AhR silencing, as it did not fully recapitulate the transcriptional shift observed in ΔAHR M-MØ (Figure 4F; Supplementary Figure 3B-D). Gene ontology analysis showed that “AhR-repressed” genes were enriched for IL-10-dependent (GSE344310) and MAFB-related genes, whereas the “AhR-dependent” module was enriched for IL-10-repressed genes (Figure 4G-H). This transcriptional shift correlated with a significant reduction in IL-10 production by FICZ-treated M-MØ (Figure 4I), consistent with direct AhR binding to the *IL10* promoter (Figure 4J). These results demonstrate that AhR restrains the anti-inflammatory program of the M-MØ subset, likely by directly repressing identity genes and limiting autocrine IL-10/MAFB signaling.

**Figure 4.**
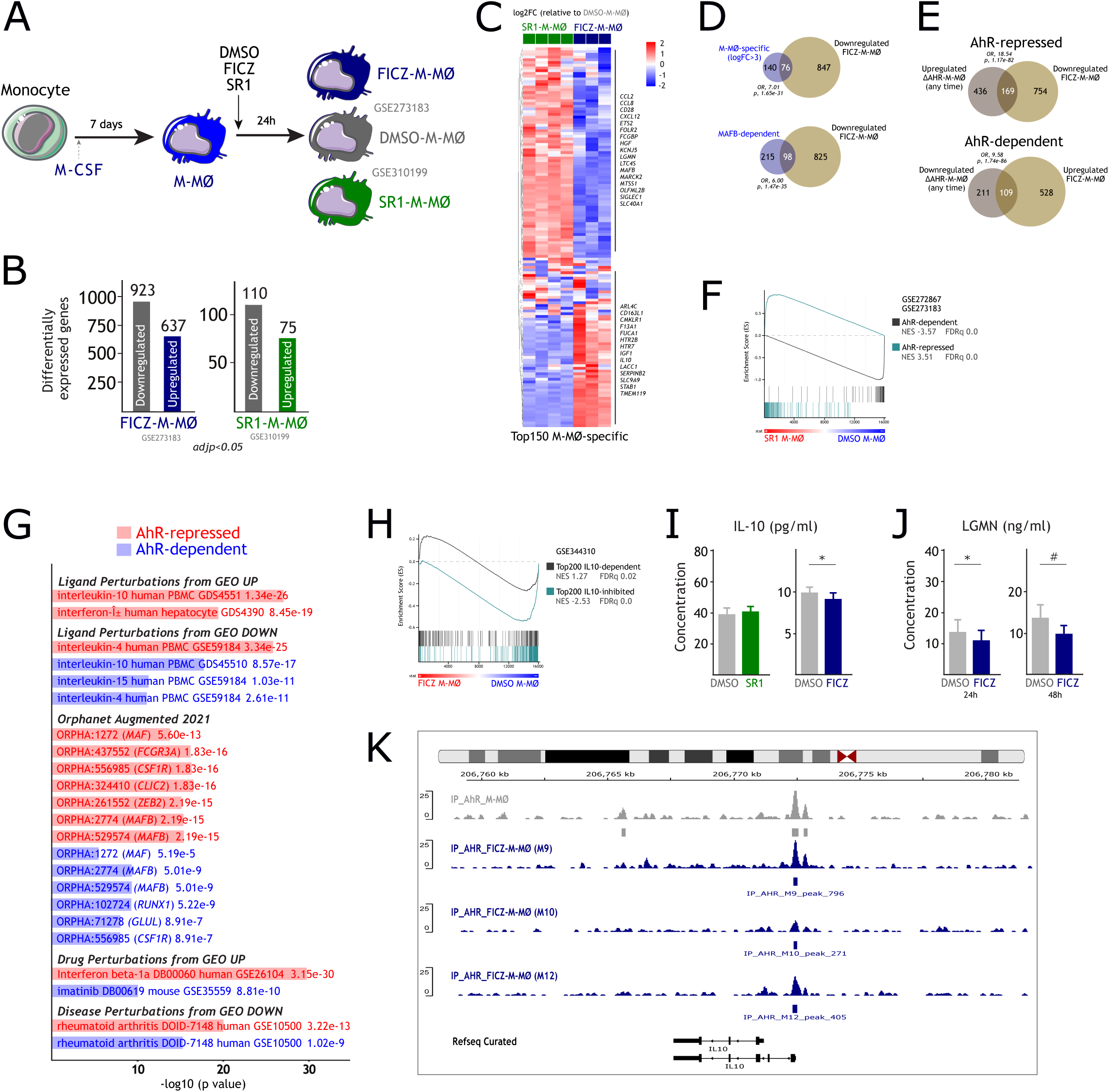
– Pharmacological modulation of AhR activity in M-MØ identifies core AhR-dependent and AhR-repressed gene sets. **A**. Generation of FICZ-, SR1-, and DMSO-treated M-MØ. **B**. Number of differentially expressed genes (adjp<0.05) in FICZ-M-MØ and SR1-M-MØ relative to DMSO-M-MØ. **C**. Heatmap of the expression of the top 150 M-MØ-specific genes in FICZ-M-MØ and SR1-M-MØ relative to DMSO-M-MØ. **D**. Overlap of genes differentially expressed in FICZ-M-MØ (relative to DMSO-M-MØ) with either M-MØ-specific genes (*Upper panel*) or MAFB-dependent genes (*Lower panel*). **E**. Intersection of genes differentially expressed in ΔAHR-M-MØ and FICZ-M-MØ to identify *bona fide* “AhR-repressed genes” (*Upper panel*) and “AhR-induced genes” (*Lower panel*). **F**. GSEA of “AhR-induced” and “AhR-repressed” gene sets on the ranked comparison of the SR1-M-MØ vs. DMSO-M-MØ gene profile. NES and FDRq are indicated. **G**. Gene ontology analysis (Enrichr) ^61^ of the “AhR-induced” and “AhR-repressed” gene sets. **H**. GSEA of the IL-10-dependent and IL-10-repressed gene sets in M-MØ (GSE344310) on the ranked comparison of FICZ-M-MØ vs. DMSO-M-MØ. NES and FDRq are indicated. **I-J**. ELISA quantification of IL-10 (**I**) and LGM (**J**) production by FICZ-M-MØ, SR1-M-MØ and DMSO-M-MØ (n=3; mean ± SEM, *, p<0.05). **K**. ChIP-Seq tracks showing AhR binding to the *IL10* gene proximal regulatory region in M-MØ.

### 5.- AhR deficiency skews M-MØ effector functions toward an exaggerated anti-inflammatory and pro-fibrotic profile

We next evaluated key M-MØ functions after AhR knockdown, specifically efferocytosis ^50^, phagocytosis, T cell activation, and activation-induced cytokine production ^51^. AhR silencing significantly enhanced efferocytosis by M-MØ (Figure 5A; Supplementary Figure 4A), aligning with upregulation of efferocytosis-related gene sets (Figure 5B; Supplementary Table S1) and consistent with an exacerbated M-MØ functional state. ΔAHR M-MØ also produced significantly higher levels of IL-10 upon LPS stimulation (Figure 5C), a trait that defines M-MØ, whereas bacterial phagocytosis remained unaffected (Figure 5D). In allogeneic co-cultures, ΔAHR M-MØ supported normal CD4+ T-cell proliferation and regulatory T lymphocyte induction (Treg; CD4^+^CD127^−^CD25^high^FOXP3^high^) (Figure 5E-F; Supplementary Figure 4B). However, responding T cells secreted higher IL-10 and lower IFN-γ levels, resulting in elevated IL-10/IFN-γ and IL-10/IL-5 ratios (Figure 5G-H), indicating that AhR limits an excessively tolerogenic T-cell priming capacity.

**Figure 5.**
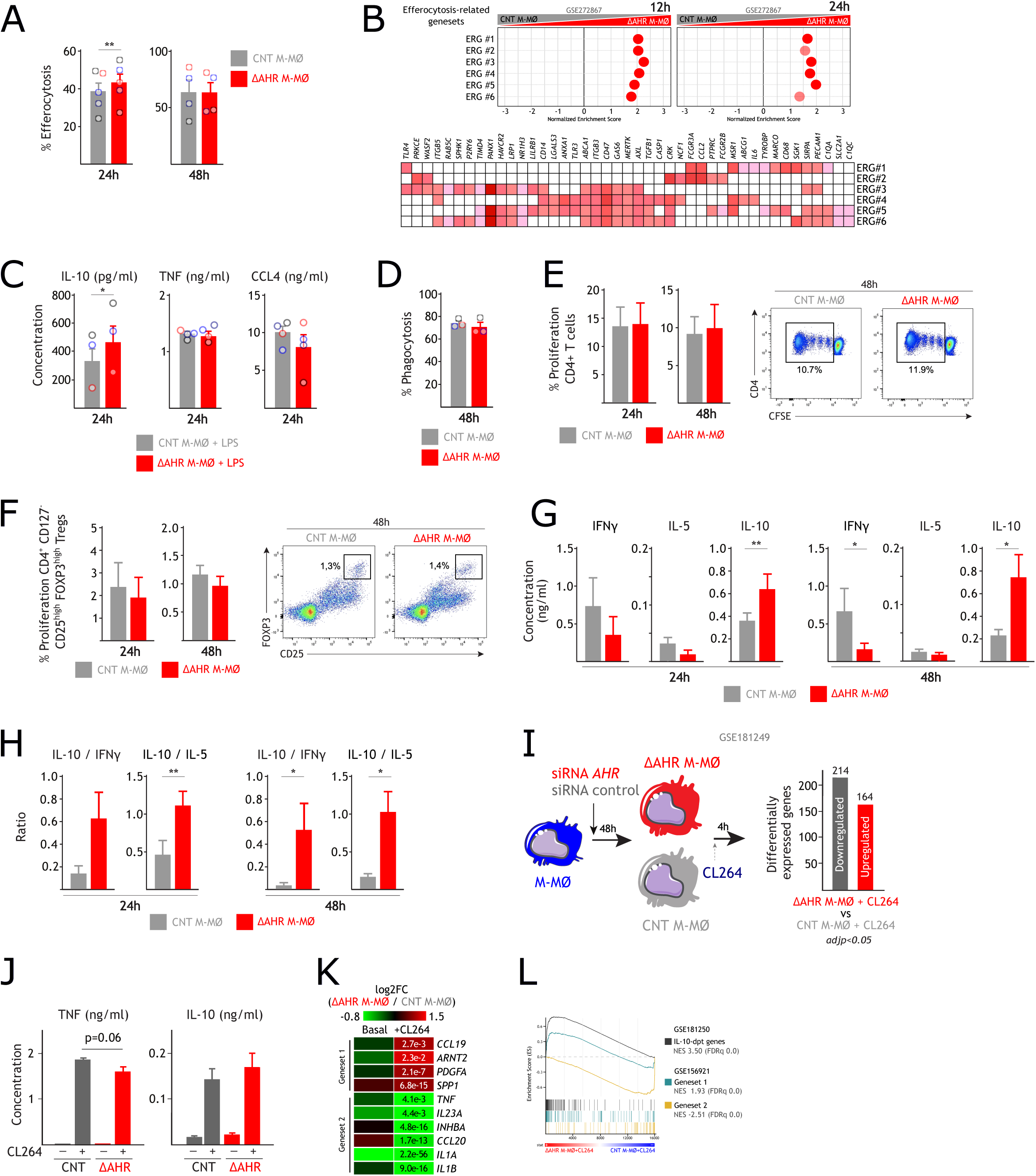
– AhR restricts efferocytosis in M-MØ and prevents exacerbated anti-inflammatory response in activated M-MØ. **A.** Efferocytosis capacity of ΔAHR-M-MØ at 24 and 48h post-transfection, showing the percentage of positive cells (n=5, 24h; n=4, 48h; mean ± SEM; paired Student’s t-test: **, p<0.01). **B**. GSEA summary (*Left panel*) and leading-edge analysis (*Right panel*) of six independent Efferocytosis-Related Gene sets (ERG#1-6) (Supplementary Table S1) on the ranked comparison of the ΔAHR-M-MØ vs. CNT M-MØ transcriptome. NES and FDRq are indicated (FDRq=0, solid color; FDR<0.25, light color) **C**. ELISA quantification of IL-10, TNF and CCL4 from LPS-treated CNT M-MØ and ΔAHR-M-MØ (n=3-5; mean ± SEM, *, p<0.05). **D.** Phagocytosis of pHRodo-labeled bacterial particles by CNT M-MØ and ΔAHR-M-MØ at 24h post-transfection, as determined by flow cytometry. The percentage of positive cells is shown (n=3; mean ± SEM). **E**. Total CD4+ T lymphocyte proliferation induced by allogeneic CNT M-MØ or ΔAHR M-MØ after 24h and 48h post-transfection, as determined by flow cytometry of CFSE-labeled cells. Mean ± SEM of 5-6 independent experiments are shown. (*Right panel*) Representative dot plots of proliferating CFSE-labelled allogeneic naive CD4+ T lymphocytes. **F**. Percentage of CD4^+^ CD127^-^ CD25^high^ FOXP3^high^ Tregs induced by allogeneic CNT M-MØ or ΔAHR M-MØ after 24h and 48h post-transfection, after 5 days of coculture (n=5-6; Mean ± SEM). Representative dot plots of induced CD4^+^ CD127^-^ CD25^high^ FOXP3^high^ Tregs are shown (*Right panel*). **G**. Cytokines produced by allogeneic naïve CD4^+^ T lymphocytes after 5 days of co-culture with allogeneic CNT M-MØ or ΔAHR M-MØ (24h and 48h post-transfection). **H**. Cytokine ratios by primed naive CD4+ T cells in the indicated conditions. **I**. Schematics for TLR7 activation (CL264) after *AHR* silencing (*Left panel*) and differential gene expression quantification between CL264-treated ΔAHR-M-MØ and CL264-treated CNT M-MØ (*Right panel*). **G**. Quantification of TNF and IL-10 released from untreated and CL264-treated ΔAHR-M-MØ and CNT M-MØ (n=4; mean ± SEM). **H**. Relative expression of selected Geneset 1 and Geneset 2 genes ^51^ in ΔAHR-M-MØ vs. CNT M-MØ, and in CL264-treated ΔAHR-M-MØ vs. CL264-treated CNT M-MØ. Statistical significance of each comparison is shown. **I**. GSEA profiles of the Geneset 1, Geneset 2, and IL-10-dependent gene sets (GSE181250), on the ranked comparison of CL264-treated ΔAHR-M-MØ vs. CL264-treated CNT M-MØ transcriptome. NES and FDRq values are indicated.

Finally, we assessed the influence of AhR deficiency on the macrophage response to pathogenic stimuli, which upregulate *AHR* expression (Supplementary Figure 4C-D). ΔAHR M-MØ stimulation with the TLR7 agonist CL264 caused a significant transcriptional divergence involving nearly 400 genes (Figure 5I), including those directly regulated by MAFB and MAF (Supplementary Figure 4E), as well as a tendency toward reduced TNF release (Figure 5J). Of note, TLR7-activated ΔAHR M-MØ showed a globally skewed activation profile, with a stronger enhancement of the characteristic M-MØ activation program (Geneset 1, including *CCL19* and *PDGFA*) (GSE156921) ^51^ and attenuated induction of the pro-inflammatory GM-MØ activation program (Geneset 2, containing *TNF*, *IL23A*, *INHBA* and *IL1A*) (Figure 5K-L). This skewed profile was significantly enriched for IL-10– and STAT3-dependent gene signatures (Figure 5L; Supplementary Figure 4F), indicating that AhR prevents over-exuberant anti-inflammatory responses during pathogen sensing.

### 6.- AhR ablation in monocytes accelerates the acquisition of a “resident-like” anti-inflammatory profile

Having established that AhR restricts the maintenance of the M-MØ anti-inflammatory profile, we next tested whether AhR governs the initial acquisition macrophage identity by ablating *AHR* in primary monocytes via CRISPR/Cas9 prior to M-CSF-driven differentiation (sgAHR M-MØ) (Figure 6A-B; Supplementary Figure 5A). sgAHR M-MØ underwent extensive transcriptional remodeling (Figure 6C), characterized by loss of AhR target signatures (Supplementary Figure 5B) and strong upregulation of core M-MØ –identity genes alongside suppression of GM-MØ markers (Figure 6D-F). GSEA confirmed enrichment for MAFB/MAF-dependent regulons and directly MAFB target genes (Figure 6G-H). This molecular switch was functionally confirmed, as sgAHR M-MØ displayed elevated basal release of CCL8, LGMN, and CXCL10, hallmark proteins of the pro-fibrotic and anti-inflammatory M-MØ profile ^52,53^ (Figure 6I), and increased LPS-induced IL-10 production (Figure 6J). Therefore, monocyte AhR ablation potentiates the acquisition of the anti-inflammatory profile, reinforcing the inverse relationship between AhR activity and the M-MØ signature. Additional gene ontology analysis revealed that the sgAHR-M-MØ transcriptome was enriched for pro-tumoral “arge TAM” ^22^ and Kupffer cells signatures, as well as MCTO-associated genes ^54^ and GSK3-inhibition profiles (Figure 6K; Supplementary Figure 5C-E). Conversely, sgAHR-M-MØ showed reduced expression of genes defining anti-tumoral “Small TAM” ^22^, infiltrating monocytes/macrophages, and downregulated upon GSK3 inhibition (Figure 6K; Supplementary Figure 5C-E). Crucially, AhR ablation markedly increased expression of ‘‘long-term resident’’ LYVE1+/ FOLR2+ macrophage (MoMac-VERSE cluster #2) ^5^ and self-renewing tissue-resident TLF^+^ (TIMD4^+^/ LYVE1^+^/FOLR2^+^) macrophage signatures ^55^ (Supplementary Figure 5F-G). Altogether, these results indicate that AhR acts as a transcriptional rheostat that restricts both the initial acquisition and the long-term maintenance of the anti-inflammatory, resident-like identity of M-CSF-driven monocyte-derived macrophages.

**Figure 6.**
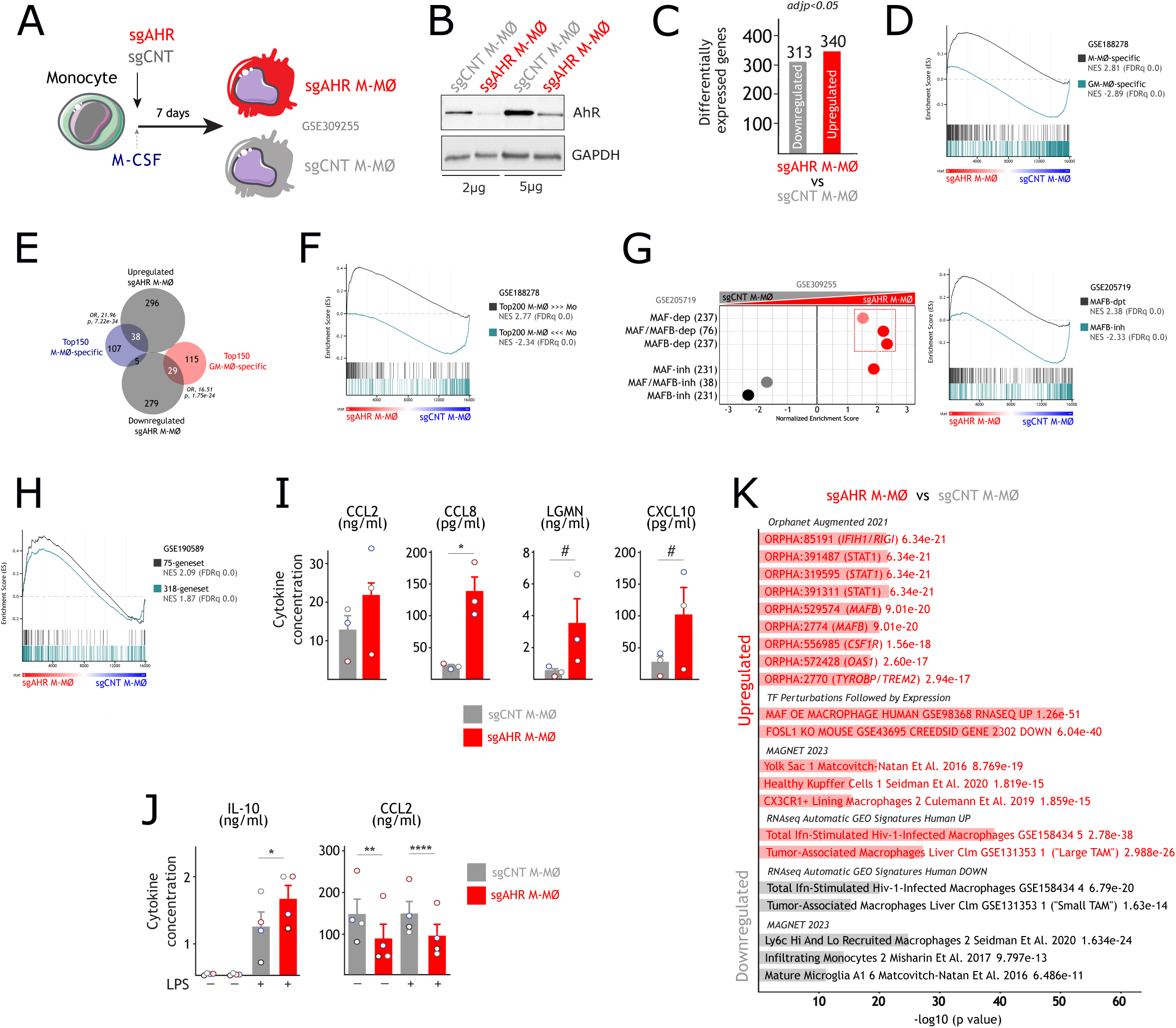
– CRISPR-mediated *AHR* deletion in monocytes restrains acquisition of the M-MØ anti-inflammatory profile. **A.** Workflow for CRISPR/Cas9 ablation of *AHR* in monocytes and subsequent differentiation into monocyte-derived M-MØ (sgAHR M-MØ). **B**. Western blot of AhR expression in sgCNT M-MØ and sgAHR M-MØ, using 2-5 μg of cell lysates. GAPDH protein levels were determined as protein loading control. **C**. Quantification of differentially expressed genes (adjp<0.05) between sgCNT M-MØ and sgAHR M-MØ. **D**. GSEA of M-MØ– or GM-MØ-specific gene sets ^67^ on the ranked comparison of the sgCNT M-MØ vs. sgAHR-M-MØ transcriptome. NES and FDRq are indicated. **E**. Overlap between genes significantly (adjp<0.05) altered in sgAHR-M-MØ (relative to sgCNT M-MØ) and the top 150 M-MØ-specific or GM-MØ-specific genes. **F**. GSEA profiles of genes differentially expressed between monocytes and monocyte-derived M-MØ ^67^ on the ranked comparison of the sgCNT M-MØ vs. sgAHR-M-MØ transcriptome. NES and FDRq values are indicated. **G**. Summary of GSEA of MAFB/MAF regulons ^41^ on the ranked comparison of the sgCNT M-MØ vs. sgAHR-M-MØ transcriptome. NES and FDRq are indicated (FDRq=0, solid colors; FDR<0.25, light colors) (*Left panel*). Two representative GSEA profiles are shown (*Right panel*). **H**. GSEA profiles of genes with MAFB-binding sites (318-geneset) and *bona fide* MAFB-dependent (75-geneset) ^21^ on the ranked comparison of the sgCNT M-MØ vs. sgAHR-M-MØ transcriptome. NES and FDRq values are shown. **I-J**. ELISA quantification of the indicated factors released by untreated (**J**) and LPS-treated (**K**) sgCNT M-MØ and sgAHR M-MØ (n=3, basal; n=4, LPS-stimulation; mean ± SEM; *, paired Student’s t-test; # ratio-paired Student’s t-test; *,^#^, p<0.05; **, p<0.01; ***, p<0.005; ****, p<0.001). **L**. Summary of the gene ontology analysis (Enrichr) ^61^ on differentially expressed genes between sgAHR-M-MØ and sgCNT M-MØ.

## DISCUSSION

The identity and functional plasticity of human macrophages are tightly regulated by a network of lineage-determining transcription factors that integrate microenvironmental signals. While the Aryl hydrocarbon receptor (AhR) is a well-known mediator of environmental sensing, its role in myeloid biology has often been simplified as a general promoter of immunosuppression. In this study, we provide a more nuanced paradigm, identifying AhR as a critical transcriptional rheostat that restrains the magnitude of the anti-inflammatory and homeostatic program in human monocyte-derived macrophages (M-MØ). By combining genomic, pharmacological, and CRISPR/Cas9 approaches, we demonstrate that AhR activity limits the expression of the MAFB/MAF-dependent regulons, thereby preventing monocyte-derived cells from adopting an unchecked pro-fibrotic and “resident-like” identity.

Although AhR is often considered an inducer of IL-10 in certain dendritic cell subsets and T regulatory (Treg) cells, where it works in concert with c-Maf to drive immunosuppression, we have found that AhR directly binds to the *IL10* proximal regulatory region and restrains its expression in M-CSF-driven MAFB-expressing macrophages. This reveals a profound cell-type-specific (context-dependent) dichotomy in AhR function. The fact that AhR ablation (via siRNA or CRISPR) significantly enhances the M-MØ signature, that is mostly driven by both MAFB and MAF transcription factors ^202137^, suggests that basal AhR activity maintains these cells in a state of functional poise. This “transcriptional brake” might prevent both an exaggerated commitment to a homeostatic phenotype and the acquisition of an exacerbated immunosuppressive state. This exacerbated tolerogenic phenotype may also influence the capacity of M-MØ to shape T-cell responses toward a more anti-inflammatory and potentially immunoregulatory profile. Given that MAFB and MAF levels remain unchanged upon AhR knockdown, this inhibitory effect might be explained by competitive binding (transcriptional interference) at shared genomic loci or antagonistic interactions between AhR and the MAF transcription factor family, effectively acting as a molecular brake on the anti-inflammatory program. Alternatively, AhR in M-MØ may recruit co-repressors/histone deacetylases to MAFB-occupied enhancers, modulating the magnitude of the anti-inflammatory response through localized chromatin remodeling. Regardless of the exact mechanism, the fact that AhR deficiency mirrors the elevated MAFB-dependent signature seen in MCTO patients highlights AhR as a critical rheostat of MAFB activity. By occupying proximal regulatory regions of M-MØ-specific genes (*e.g*., *CD163*, *LGMN*), AhR may obstruct or diminish the recruitment of the transcriptional machinery, ensuring that the anti-inflammatory profile is maintained within homeostatic limits.

The repression of IL-10 by AhR appears to be a central node in this process. Since IL-10 is a potent autocrine inducer of MAFB expression and STAT3 phosphorylation, AhR-mediated repression of IL10 might serve as a dual-layered barrier: it directly limits the transcription of identity genes and indirectly inhibits the signaling axis that sustains them. When AhR is absent, this barrier is removed, leading to an overactive IL-10/STAT3/MAFB axis that functionalizes the cells toward pro-fibrotic/repair signature, as evidenced in our TLR7 stimulation experiments, where ΔAHR M-MØ failed to mount a proper pro-inflammatory response and instead skewed toward a pro-fibrotic repair signature (Geneset 1). Our pharmacological data further support this model, as the agonist FICZ dampens the M-MØ profile, while SR1-mediated AhR inhibition partially mirrors the effects of AhR silencing. This suggests that AhR may exert its effects not only through ligand-induced activity but potentially through ligand-independent basal occupancy or steric hindrance.

An additional significant finding in our study is that AhR might act as a molecular gatekeeper of macrophage ontogeny. Tissue-resident macrophages (like Kupffer cells or alveolar macrophages) are largely defined by their embryonic origin and self-renewal capacity, as opposed to infiltrating monocyte-derived macrophages. However, our CRISPR/Cas9 model reveals that monocytes lacking AhR acquire the molecular hallmarks of “long-term resident” TLF^+^ macrophages and the MoMac-VERSE cluster #2 subset, populations typically associated with minimal monocyte contribution. This suggests that monocyte-derived macrophages possess a latent plastic potential to adopt a resident-like identity that is actively restrained by AhR. By acting as a transcriptional barrier, AhR might therefore ensure that monocyte-derived cells remain functionally plastic and distinct from niche-specialized, self-renewing resident populations. Thus, AhR would ensure that monocyte-derived macrophages remain poised for transient responses rather than achieving the profound homeostatic commitment of tissue-resident TLF+ macrophages. Therefore, AhR might be a key regulator of the “monocyte-to-resident” transition, and the modulation of the AhR-MAFB axis may provide a novel therapeutic strategy to ‘re-program’ monocyte-derived cells into homeostatic, resident-like states in the context of chronic inflammation or fibrosis.

The clinical relevance of the AhR-regulated gene signature is underscored by its enrichment in various disease-associated macrophage subsets. We found that AhR deficiency enhances signatures associated with pro-fibrotic macrophages in severe COVID-19 and pro-tumoral TAM in colorectal liver metastasis. In these contexts, macrophages often adopt a hyper-anti-inflammatory profile that promotes tissue remodeling and immunosuppression at the expense of host defense. Our data suggest that the endogenous upregulation of AhR during activation (such as TLR7 sensing) serves as a necessary “brake” to resolve inflammation and prevent fibrosis. However, in chronic diseases, this mechanism may be subverted. Therapeutically, hyper-activation of AhR could be leveraged to curb the expansion of pathogenic, pro-fibrotic macrophages. Conversely, in oncology, AhR antagonists might be used to prevent TAM from adopting a “resident-like” immunosuppressive state, potentially increasing susceptibility to immunotherapy, although AhR activation in tumor-associated macrophages by microbial metabolites has been shown to suppress anti-tumor immunity ^56^. The transition from anti-tumoral “Small TAM” to pro-tumoral “Large TAM” often marks a shift towards immunosuppression and metastasis. By limiting the magnitude of the MAFB-driven anti-inflammatory program, AhR might ensure that macrophages maintain a degree of plasticity. We propose that the AhR-MAFB axis represents a critical checkpoint in macrophage ontogeny; while MAFB drives the acquisition of the M-MØ identity, AhR fine-tunes this process to prevent the hyper-activation of pathogenic gene modules.

In conclusion, we have identified AhR as a transcriptional rheostat that defines the functional boundaries of human macrophages. By limiting the IL-10/MAFB axis, AhR appears to prevent monocyte-derived macrophages from excessively committing to a homeostatic or pro-fibrotic state. This molecular checkpoint maintains the balance between inflammatory responsiveness and tissue repair, positioning AhR as a promising target for the reprogramming of macrophage-mediated pathologies.

## MATERIALS AND METHODS

### Generation of monocyte-derived macrophages and ELISA

Buffy coats from anonymous healthy blood donors were provided by Comunidad de Madrid blood Bank. Ethical approvals for all blood processes used in this study were approved by the Centro de Investigaciones Biológicas Ethics Committee, and all experiments were carried out in accordance with the approved guidelines and regulations. Human peripheral blood mononuclear cells (PBMC) were isolated from buffy coats over a Lymphoprep (Nycomed Pharma) gradient, and monocytes were purified from PBMC by magnetic cell sorting using CD14 microbeads (Miltenyi Biotech). Monocytes (>95% CD14+ cells) were cultured at 0.5 x 10^6^ cells/ml for 7 days in RPMI 1640 supplemented with 10% fetal bovine serum (FBS, Gibco) (complete medium) and M-CSF (10 ng/ml) to generate M-MØ or GM-CSF (1000 U/mL) to generate GM-MØ, as previously described ^19^. For macrophage activation, cells were treated with 10 ng/ml E. coli 055:B5 lipopolysaccharide (LPS, Sigma-Aldrich, MO, USA). Where indicated, macrophages were treated with the AhR antagonists CH223191 (1-methyl-*N*-[2-methyl-4-[2-(2-methylphenyl)diazenyl]phenyl]-*1H*-pyrazole-5-carboxamide, 3 µM) or SR1 (StemRegenin1, 0.5 μM), the AhR ligand FICZ (6-formylindolo[3,2-b]carbazole, 0.25 μM) or DMSO as a control. Cytokine detection was carried out with ELISA kits for human CCL2 (BD Biosciences, #555179), CCL8 (R&D, #DY281), Legumain (LGMN) (R&D, #DY4769), IL-10 (R&D, #DY217B), CXCL10 (BioLegend, #439904) and TNF (BD Biosciences, #555212).

### Quantitative PCR

Total RNA was extracted using the total RNA and protein isolation kit (Macherey-Nagel). RNA samples were reverse transcribed with High-Capacity cDNA Reverse Transcription reagents kit (Applied Biosystems) according to the manufacturer’s protocol. Quantitative PCR (qPCR) was performed with Light-Cycler 480 Probes Master (Roche Life Sciences) and TaqMan probes on a standard plate in a Light Cycler 480 instrument (Roche Diagnostics). Gene-specific oligonucleotides were designed using the Universal ProbeLibrary software (Roche Life Sciences). Results were normalized to the expression level of the endogenous reference genes *TBP* and *HPRT1* and quantified using the ΔΔCT method.

### RNA-Seq and data analysis

RNA was isolated from M-MØ transfected with *AHR*-specific siRNA (ΔAHR M-MØ), M-MØ treated with FICZ (FICZ-M-MØ), M-MØ treated with SR1 (SR1-M-MØ), GM-MØ transfected with *AHR*-specific siRNA (ΔAHR GM-MØ), and M-MØ derived from monocytes transfected with sgAHR (sgAHR M-MØ), and subjected to sequencing on a BGISEQ-500 platform (http://www.bgitechsolutions.com/). Low-quality reads and reads with adaptors or unknown bases were filtered to get the clean reads. Sequences were mapped to GRCh38 genome using HISAT2 ^57^ or Bowtie2 ^58^, and clean reads for each gene were calculated using htseq-count ^59^ and the RSEM software package ^60^. RNA-Seq data were deposited in the Gene Expression Omnibus (http://www.ncbi.nlm.nih.gov/geo/) under accession no. GSE272867 (ΔAHR M-MØ), GSE273183 (FICZ-M-MØ), GSE310199 (SR1-M-MØ), GSE309272 (ΔAHR GM-MØ) and GSE309255 (sgAHR M-MØ). Differential gene expression was assessed by using the R-package DESeq2, and analyzed for annotated gene sets enrichment using ENRICHR (http://amp.pharm.mssm.edu/Enrichr/) ^61^, and enrichment terms considered significant with a Benjamini-Hochberg adjp<0.05. For gene set enrichment analysis (GSEA) (http://software.broadinstitute.org/gsea/index.jsp) ^62^, gene sets available at the website, as well as gene sets generated from publicly available transcriptional studies (https://www.ncbi.nlm.nih.gov/gds), were used. AhR-dependent gene sets were extracted from previous studies reporting: 1) genes positively or negatively regulated by BaP in human macrophages ^32^; 2) genes regulated by VAF347 in human dendritic cells ^33^ (GSE10463); 3) genes regulated by I3C in murine macrophages ^34^ (GSE223122); 4) genes regulated by SR1 in AML cells ^35^ (GSE48843); and 5) genes specifically bound by AhR in ChIP-Seq on MCF-7 cells ^36^. Additional data sets used throughout the manuscript (either reported here or previously published by our group or others) are listed and described in Supplementary Table S1.

### Western blot

Cell lysates were subjected to SDS-PAGE (50 μg unless indicated otherwise) and transferred onto an Immobilon-P polyvinylidene difluoride membrane (PVDF; Millipore, Sigma). After blocking the unoccupied sites with 5% nonfat milk diluted in Tris-Buffered Saline plus Tween 20 (TBS-T), protein detection was carried out with antibodies against AhR (D5S6H, Cell Signaling; sc-133088, Santa Cruz Biotechnology), CD163 (MCA1853, BIO-RAD), Lamin B1 (D4Q47, Cell Signaling), MAFB (HPA005653, Sigma-Aldrich), and MAF (E2Q5D, Cell Signaling), or GAPDH (sc-32233, Santa Cruz Biotechnology) and Vinculin (V9131, Sigma-Aldrich) as a protein loading control. Chemiluminescence was detected in a Chemidoc Imaging system (Bio-Rad) using SuperSignal West Femto (Thermo Fisher Scientific).

### siRNA transfection

Monocyte-derived macrophages (1 × 10^6^ cells) were transfected with a human *AHR*-specific siRNA (siAHR, 50 nM) (Ambion) using HiPerFect (Qiagen). Silencer Select Negative Control No. 1 siRNA (siCtrl, 50 nM) (Ambion) was used as negative control siRNA. Cells and cell supernatants were collected 12/24/48 hours after transfection and processed.

### AHR knockdown in primary human monocytes using CRISPR-Cas9 ribonucleoprotein complex

Human primary human monocytes were isolated from peripheral blood mononuclear cells from buffy coats using a Lymphoprep (Nycomed Pharma) gradient and subsequently purified by magnetic cell sorting using anti-CD14 microbeads (Miltenyi Biotec) (>95% CD14+ cells). To assemble ribonucleoprotein (RNP) complexes, AHR sgRNA (multi-guide RNA strategy, Gene Knockout Kit v2 – human – AHR, Synthego) or negative control sgRNA (Negative Control sgRNA #1 Synthego) were resuspended in sterile water at a concentration of 100 μM, and RNP complexes generated by incubating 500 pmol of AHR sgRNA (or negative control sgRNA) with 200 pmol of Cas9 (stock of 20 μM Cas9 Nuclease 2NLS, *S. pyogenes*, Synthego) at 37°C for 15-20 minutes. Nucleofection of Cas9-RNP complexes into primary human monocytes was done according to established procedures (protocol for Primary Human Monocytes for 4D-Nucleofectort X-Unit, Lonza). In brief, 8×10^6^ monocytes were centrifugated (100*g*, 9 minutes), carefully resuspended in 4D nucleofector solution (P3+ Supplement, Lonza) at room temperature, mixed with 15 μl of the RNP mixtures, transferred into 100 μl nucleocuvettes, and nucleofected in the 4D –Nucleofector X Unit (Lonza) using the EA-100 program. Immediately after nucleofection, 500 μl of pre –warmed complete RPMI 1640 (containing 10% fetal bovine serum and 10 ng/ml M-CSF, ImmunoTools) was added to the cuvettes and maintained in the dark at 37°C 5% CO2 for 15-30 minutes. After careful mixing, 2.6×10^6^ monocytes were added into 12-well plates containing 2 ml of pre-warmed complete medium. Then, cells were incubated following established protocols to generate M-CSF-dependent monocyte-derived macrophages (M-MØ). To generate cell lysates, each well containing nucleofected cells was washed with PBS, lysed using 30 μl of 5% SDS and incubated for 3 minutes at room temperature. Cell lysates (90 μl /nucleofection) were transferred to 1.5 ml Eppendorf tubes and heated to 95°C for 3 minutes. After centrifugation (12000 rpm, 1 min), supernatants were recovered and stored at –20°C.

### Chromatin immunoprecipitation and sequencing (ChIP-Seq)

ChIP-Seq was performed essentially as described ^6321^. M-MØ generated from two independent donors (150×10^6^ cells) were combined, pelleted and fixed with 1% formaldehyde in PBS for 15 minutes. Fixation was stopped using glycine 0.2M. After centrifuging (8 min at 3000 rpm, 4°C), cells were washed twice with cold PBS by centrifugation (8 min at 3000 rpm, 4°C). After supernatant removal, pellet was lysed in hypotonic buffer containing protease inhibitors (10 minutes, 4°C, 300 rpm), and this step was repeated once more. Lysate was homogenized and stored at –80°C before sonication. Sonication was done in a Diagenode Bioruptor UCD-200TM-EX, with 30 cycles of 30“ON-30”OFF, with vortexing every 10 cycles. Sonicated lysate was then centrifuged at 11000 rpm for 10 minutes at 15°C. 10% of the supernatant was used as INPUT and kept at –80°C. De-crosslink was done for 4.5 hours at 65°C using 250mM NaCl and 400 μg/ml of proteinase K. Input DNA was purified using the QIAquick PCR purification kit from Qiagen. DNA fragmentation was assessed with the High Sensitivity Bioanalyzer Agilent 2100. For chromatin immunoprecipitation, preclearing and immunoprecipitation were done using the anti-AhR rabbit monoclonal antibody D5S6H (Cell Signaling Technology, #83200), and using a rabbit polyclonal IgG as a control. Washing steps were done following standard procedures (3 washes with a low-salt buffer, 3 washes with a high-salt buffer, 3 washes with a LiCl-containing buffer). After de-crosslinking with 250 mM NaCl (2 hours at 65°C, with agitation at 350 rpm), DNA was treated with 400 micrograms/ml of proteinase K (1 hour at 37°C). Then, DNA was purified using the IPure kit v2 from Diagenode. Sequencing of ChIP-Seq–derived libraries was performed on the BGI-500 platform. Sequenced single-end 50 bp reads were aligned to the genome assembly GRCh38. MACS3 (https://github.com/macs3-project/MACS) ^64^, ChIPseeker ^65^ and MEME (https://github.com/cinquin/MEME) ^66^ were used for peak calling, peak annotation and motif discovery, respectively. The Integrative Genomics Viewer (IGV) genome browser was used to visualize the aligned read files and the identified peaks (https://software.broadinstitute.org/software/igv/). ChIP-Seq data was deposited in GEO under accession no. GSE273002 (untreated M-MØ) and GSE273003 (FICZ-treated M-MØ).

### Efferocytosis and Phagocytosis assays

For efferocytosis, Jurkat T cells were rendered apoptotic by Staurosporine treatment (0.5 μg/ml) for 3 h at 37 °C (population of 83% annexin V + cells), and Apoptotic cells (AC, 1 × 10^6^ cells/ml) were labeled with CellTrace™ Violet (0.5 μM, Invitrogen) for 20 min. Macrophages were co-incubated with labeled AC (ratio 1:41; h at 37 °C) in 24-well plates, washed thoroughly with PBS to remove unbound AC and detached with PBS 5 mM EDTA, and pelleted by centrifugation before analyzing by flow cytometry. For Phagocytosis, macrophages were incubated with pHrodo Red *E. coli* BioParticles Conjugates (Thermo Fisher) for 60 min in 24-well plates at 37 °C and with 5% CO_2_, harvested and analyzed by flow cytometry.

### Allogeneic Mixed Lymphocyte Reaction

Naïve CD4+ T cells were isolated from human PBMCs (from donors different from those used for macrophage generation) using the Naïve CD4+ T Cell Isolation Kit (Miltenyi Biotec) in an autoMACS Pro separator (Miltenyi Biotec), according to the manufacturer’s instructions. Purified naïve CD4+ T cells were subsequently labeled with 5 µM CFSE (Invitrogen). Macrophages were detached using PBS-EDTA, collected, counted and resuspended in complete RPMI 1640 medium [RPMI 1640, Lonza, supplemented with 10% heat-inactivated fetal bovine serum (FBS, HyClone), 100 µg/mL Normocin (InvivoGen), 50 µg/mL penicillin-streptomycin, 1% nonessential amino acids, 1% MEM vitamins and 1 mM sodium pyruvate (all from Life Technologies)]. For MLR, 1 × 10^5^ macrophages were cocultured with 5 × 10^5^ naïve CD4+ T cells (1:5 macrophage-to-T-cell ratio) in 48-well plates (500 µL per well). Cocultures were maintained for 5 days at 37°C in a humidified atmosphere containing 5% CO₂. Following the 5-day coculture, cells and supernatants were collected for downstream analyses. Cells were stained for viability using LIVE/DEAD Fixable Aqua (Invitrogen), followed by surface staining with BB700 anti-human CD4 (BD Biosciences), BV421 anti-human CD25 (BioLegend), and PE-labeled anti-human CD127 (BioLegend). Cells were subsequently fixed and permeabilized using FOXP3 Fix/Perm and Perm/Wash Buffers (BioLegend) and stained intracellularly with Alexa Fluor 647-labeled anti-human FOXP3 (BioLegend). Flow cytometry analysis was performed using a FACSymphony A1 cytometer (BD Biosciences) in the Flow Cytometry and Fluorescence Microscopy Unit at Complutense University of Madrid to assess T-cell proliferation based on CFSE dilution, and the induction of CD4^+^ CD127^−^ CD25^high^ FOXP3^high^ regulatory T-cells (Treg). Flow cytometry data were analysed using FlowJo V10 software.

### Reporter gene assay

Reporter gene assays were performed essentially as described ^29^. The Cignal Xenobiotic Response Element (XRE) Reporter Assay Kit (Qiagen), in which the XRE-firefly luciferase construct harbors tandem repeats of the AhR-binding element (XRE), was used for the reporter gene assays. For normalizing transfection efficiency, transfected DNA included a 40:1 mixture of the XRE-specific firefly luciferase construct and a construct expressing *Renilla* luciferase from a constitutive promoter. Cells (1×10^6^ per well) were transfected with 1 μg of the XRE reporter construct using jetOPTIMUS (Polyplus). After transfection, cells were cultured overnight before exposure to FICZ (250 nM), CH223191 (3 μM), or DMSO for 24 h and lysed. Firefly and *Renilla* luciferase activities were determined using the Dual-Luciferase Reporter Assay System (Promega).

### Statistics

Statistical analyses were conducted using the GraphPad Prism software. For comparison of means, and unless otherwise indicated, statistical significance of the generated data was evaluated using repeated-measures one-way ANOVA followed by Dunnett’s multiple-comparison test, paired Student’s t-test (2-tailed; *) or paired ratio Student’s t-test (2-tailed; #). In all cases, p < 0.05 was considered statistically significant.

### Data availability

The data set supporting the conclusions of this article is available in the Gene Expression Omnibus repository (http://www.ncbi.nlm.nih.gov/geo/) under accessions GSE272867 (ΔAHR M-MØ), GSE273183 (FICZ-M-MØ), GSE310199 (SR1-M-MØ), GSE309272 (ΔAHR GM-MØ) and GSE309255 (sgAHR M-MØ), GSE273002 (ChIP-Seq on untreated M-MØ) and GSE273003 (ChIP-Seq on FICZ-treated M-MØ).

## ACKNOWLEDGEMENTS

This work was supported by: grants PID2020-114323RB-I00 and PID2024-157082OB-I00 (to ALC), grant PID2024-159113OB-I00 (to AS), and grant PID2024-156862OB-I00 (to OP) from Ministerio de Ciencia, Innovación y Universidades/Agencia Estatal de investigación (MICIU/AEI), and co-funded by “FEDER: A way to make Europe”; and grant P2022/BMD-7274 (to ALC) from Dirección General de Innovación e Investigación Tecnológica de la Comunidad de Madrid. ASM was funded by grant AGAUR_FI23-2023FI_1_00771 from the Generalitat de Catalunya. ACB, IR and MSF were funded by a Formación de Personal Investigador predoctoral fellowships from Ministerio de Ciencia e Innovación (Grants PRE2022-101330, PRE2021-097080 and PRE2018-083396, respectively).

