## Supplementary Figures and table for "Aryl hydrocarbon receptor (AhR) restrains the maintenance and acquisition of the anti-inflammatory identity of monocyte-derived macrophages (M-MØ)"

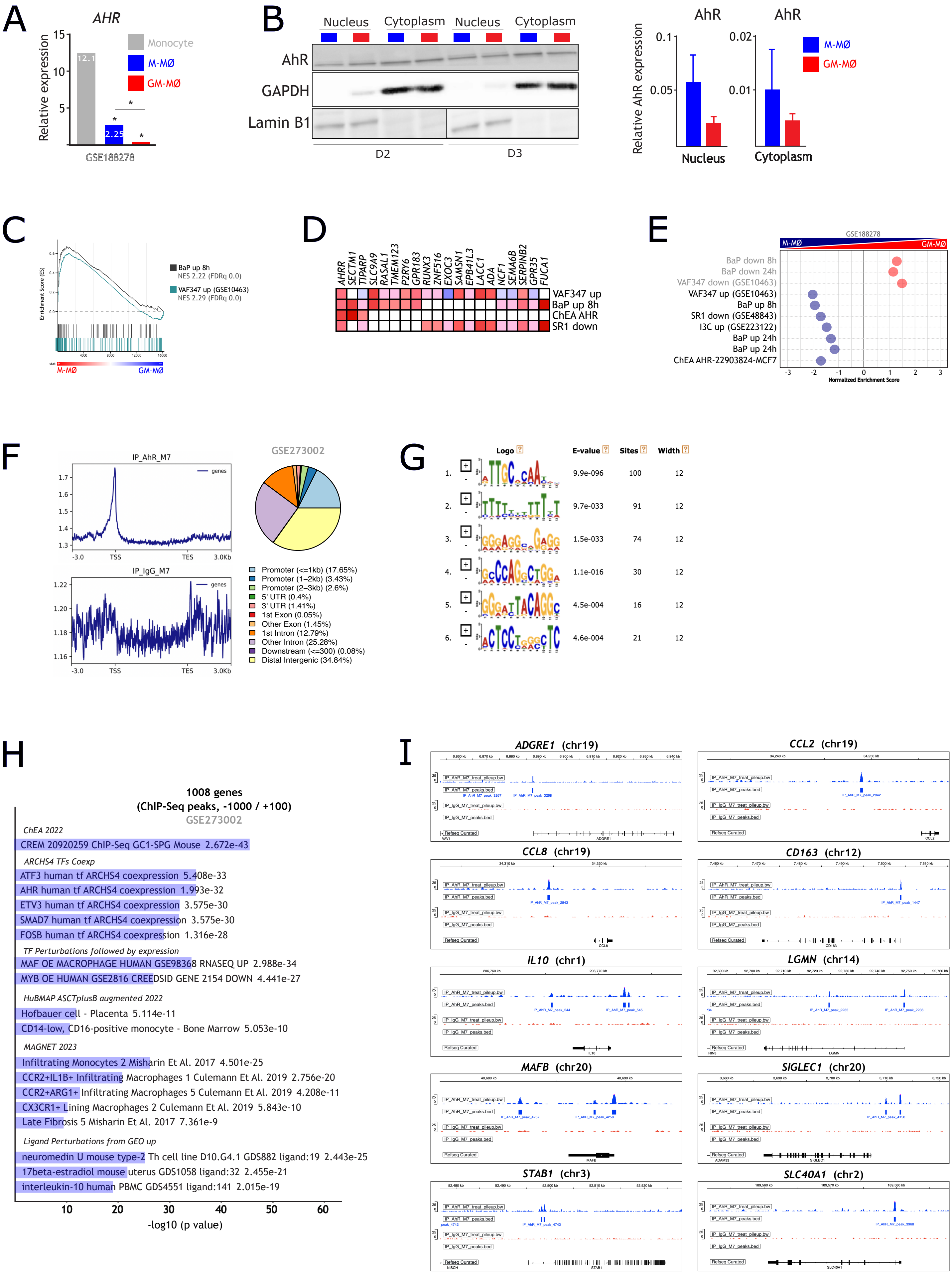


Supplementary Figure 1

**Supplementary Figure 1.- AhR expression, localization, and genomic binding in human macrophages**. **A**. Relative *AHR* mRNA expression in monocytes and monocyte-derived M-MØ and GM-MØ macrophages [3839] (*, adjp<0.05). **B**. Subcellular distribution of AhR in M-MØ and GM-MØ (n=2) determined by Western blot (*Left panel*). GAPDH and Lamin B1 served as cytoplasmic and nuclear markers, respectively. Mean ± SEM of the AhR/GAPDH and AhR/Lamin B1 protein ratios are shown (*Right panel*). **C**. GSEA profiles of the genes upregulated by VAF347 in dendritic cells [33], or BaP in GM-MØ [32], on the ranked comparison of the GM-MØ vs. M-MØ transcriptome. NES and FDRq values are shown. **D**. Leading-edge analysis for GSEA results shown in panel **D**. **E**. GSEA summary of canonical AhR-regulated genes sets [from [3233343536]] on the ranked comparison of the GM-MØ vs. M-MØ transcriptome from GSE188278 [39] NES and FDRq values are shown (light colors, FDRq<0.25). **F**. Average ChIP-Seq signal intensity for an anti-AhR rabbit monoclonal antibody D5S6H (Cell Signaling Technology, #83200) (IP_AhR_M7) or a rabbit polyclonal IgG (IP_IgG_M7) across a normalized gene body (TSS, Transcriptional Start Site; TES, Transcriptional End Site) (GSE273002). The relative genomic locations of the ChIP-Seq AhR peaks is shown. **G**. Motif enrichment within ChIP-Seq AhR peaks, with indication of position weight matrices and statistical significance. **H**. Enrichr gene ontology analysis of the 1008 genes with AhR occupancy at proximal regulatory regions (-1000/+100). **I**. Representative ChIP-Seq tracks showing AhR binding sites at specific M-MØ identity genes.

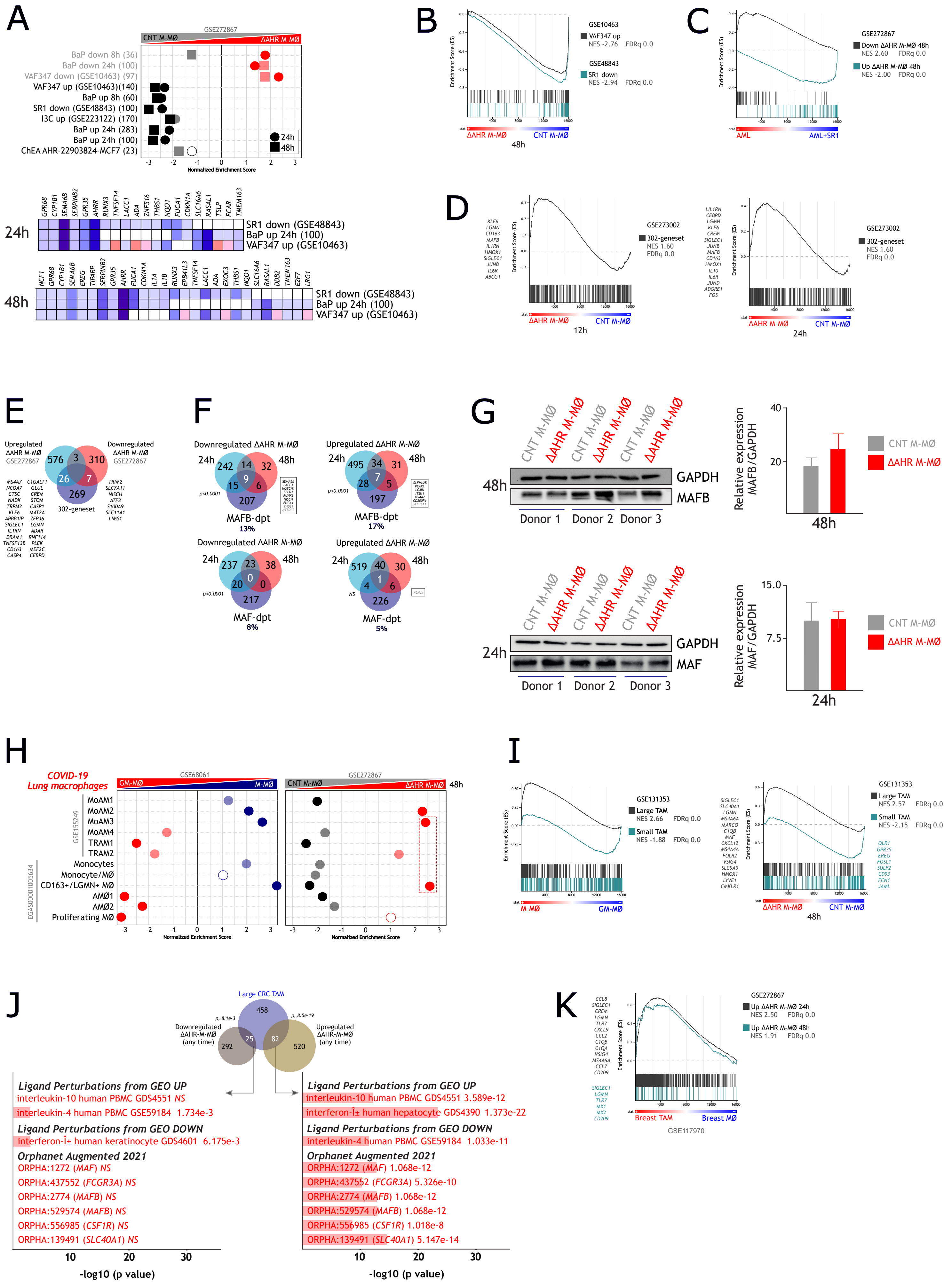


Supplementary Figure 2

**Supplementary Figure 2.- AhR silencing potentiates the M-MØ signature**. **A**. GSEA and leading-edge analysis of canonical AhR signatures [from [3233343536]] on the ranked comparison of the CNT M-MØ vs. ΔAHR M-MØ transcriptome at 24h and 48 after siRNA transfection. NES and FDRq values are indicated (FDRq=0, solid colors; FDR<0.25, light colors; Not significant, empty circles). **B**. GSEA profiles of the genes downregulated by SR1, or upregulated by VAF347, on the ranked comparison of the CNT M-MØ vs. ΔAHR M-MØ transcriptome at 48 after siRNA transfection. NES and FDRq values are indicated. **C**. GSEA profiles of the genes specifically up- or downregulated in ΔAHR M-MØ 48 after siRNA transfection on the ranked comparison of the transcriptomes of untreated Acute Myeloid Leukemia (AML) vs. SR1-treated AML (AML+SR1). NES and FDRq values are indicated. **D**. GSEA profile of the 302-gene AhR cistrome (high occupancy ChIP-Seq peaks) on the ranked comparison of the CNT M-MØ vs. ΔAHR M-MØ transcriptome at 12h and 24h after siRNA transfection. NES and FDRq values are indicated, and representative leading-edge genes are shown. **E**. Overlap between genes differentially expressed in ΔAHR-M-MØ and the 302-gene ChIP-Seq cistrome. **F**. Overlap of genes differentially expressed in ΔAHR-M-MØ and the MAFB/MAF-dependent regulons. **G**. Western blot and quantification (MAFB/GAPDH and MAF/GAPDH ratios) of MAFB and MAF protein levels in CNT M-MØ and ΔAHR-M-MØ (n=3; mean ± SEM). **H**. GSEA of the gene sets that characterize the macrophage subsets identified in severe COVID-19 [4415] on the ranked comparison of the GM-MØ vs. M-MØ transcriptome (*Left panel*) or the CNT M-MØ vs. the ΔAHR-M-MØ transcriptome at 48h after transfection (*Right panel*). NES and FDRq values are indicated (FDRq=0, solid colors; FDR<0.25, light colors; Not significant, empty circles). **I**. GSEA profiles of the gene sets that define "Large TAM" and "Small TAM" subsets from colorectal liver metastasis (GSE131353) [22] on the ranked comparison of the GM-MØ vs. M-MØ transcriptome (GSE68061) (*Left panel*) or the CNT M-MØ vs. ΔAHR-M-MØ transcriptome at 48h after transfection (GSE272867) (*Right panel*). Representative leading-edge genes are shown. **J.** Overlap of ΔAHR-M-MØ modulated genes (adjp<0.05 at any time point) and genes within the "Large TAM" subset from colorectal liver metastasis (GSE131353) [22]. Gene ontology analysis (Enrichr) [61] of the most significant overlap is shown. **K**. GSEA profiles of the genes upregulated in the ΔAHR-M-MØ transcriptome at 24 or 48h after transfection on the ranked comparison of the Breast MØ vs. Breast TAM transcriptome. Representative leading-edge genes are shown.

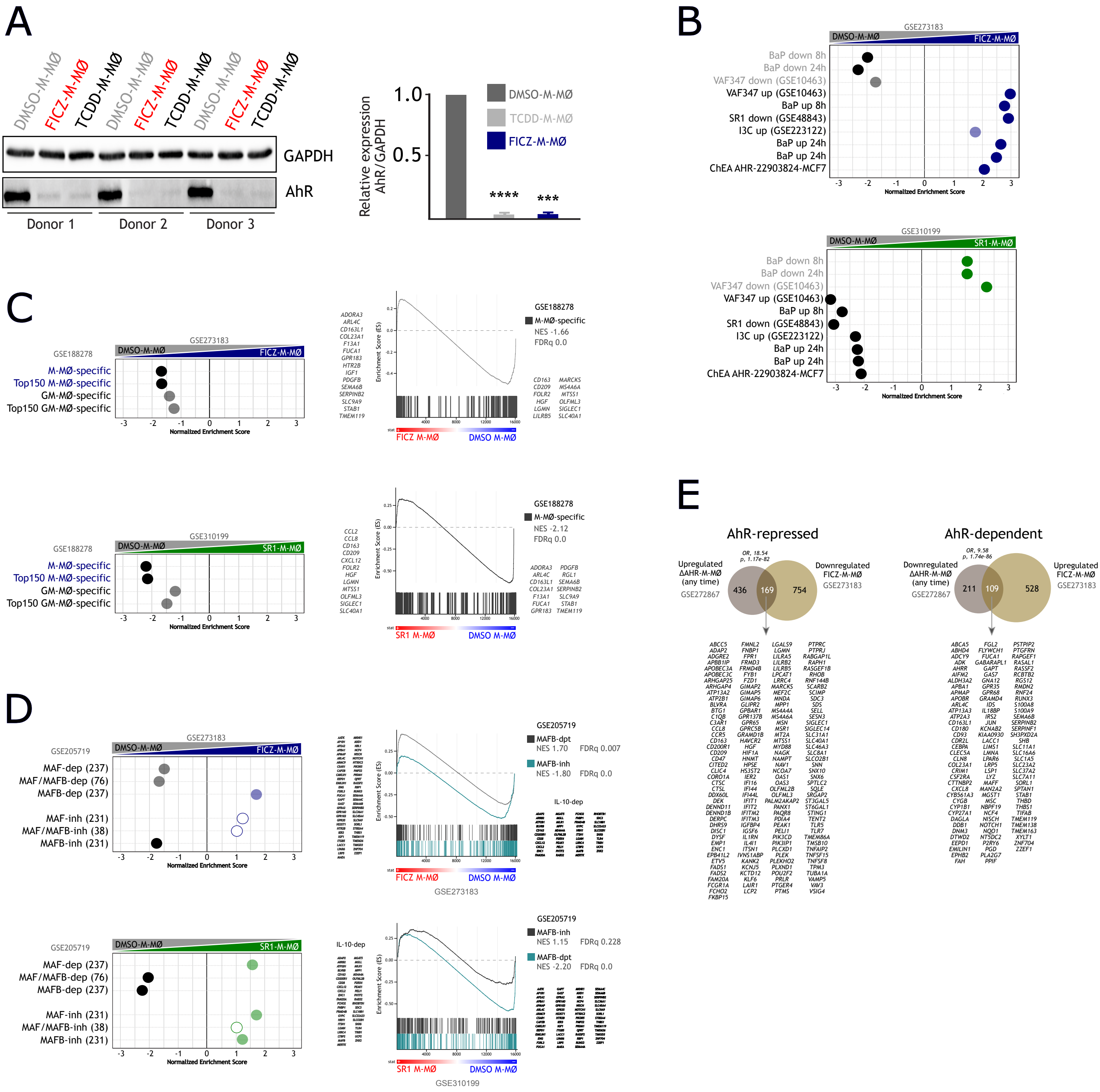


Supplementary Figure 3

**Supplementary Figure 3.- Pharmacological modulation of AhR activity shifts the macrophage identity program. A.** Western blot (*Left panel*) and quantification (AhR/GAPDH ratio, *Right panel*)) of AhR in M-MØ treated with FICZ, TCDD or DMSO (n=3; mean ± SEM; ****, $p<0.001$). **B**. GSEA summary of canonical AhR targets [from [3233343536]] on the ranked comparison of either the FICZ-M-MØ vs. DMSO-M-MØ transcriptome (*Left panel*) or the SR1-M-MØ vs. DMSO-M-MØ transcriptome (*Right panel*). NES and FDRq values are indicated (FDRq=0, solid colors; FDRq<0.25, light colors). **C**. GSEA summary (*Left panels*) and representative profiles (*Right panels*) of the indicated M-MØ- or GM-MØ-specific gene sets [67] on the ranked comparison of the FICZ-M-MØ vs. DMSO-M-MØ transcriptome (*Upper panels*) or SR1-M-MØ vs. DMSO-M-MØ transcriptome (*Lower panels*). NES and FDRq values are indicated (FDRq=0, solid colors; FDRq<0.25, light colors). Representative leading-edge genes are shown. **D**. GSEA summary (*Left panels*) and representative profiles (*Right panels*) of MAFB/MAF regulons [41] on the ranked comparison of either the FICZ-M-MØ vs. DMSO-M-MØ transcriptome (*Upper panels*) or SR1-M-MØ vs. DMSO-M-MØ transcriptome (*Lower panels*). NES and FDRq values are indicated (FDRq=0, solid colors; FDR<0.25, light colors; Not significant, empty circles). Representative leading-edge genes are shown. **E**. Identification of the *bona fide* "AhR-induced" and "AhR-repressed" genes in M-MØ.

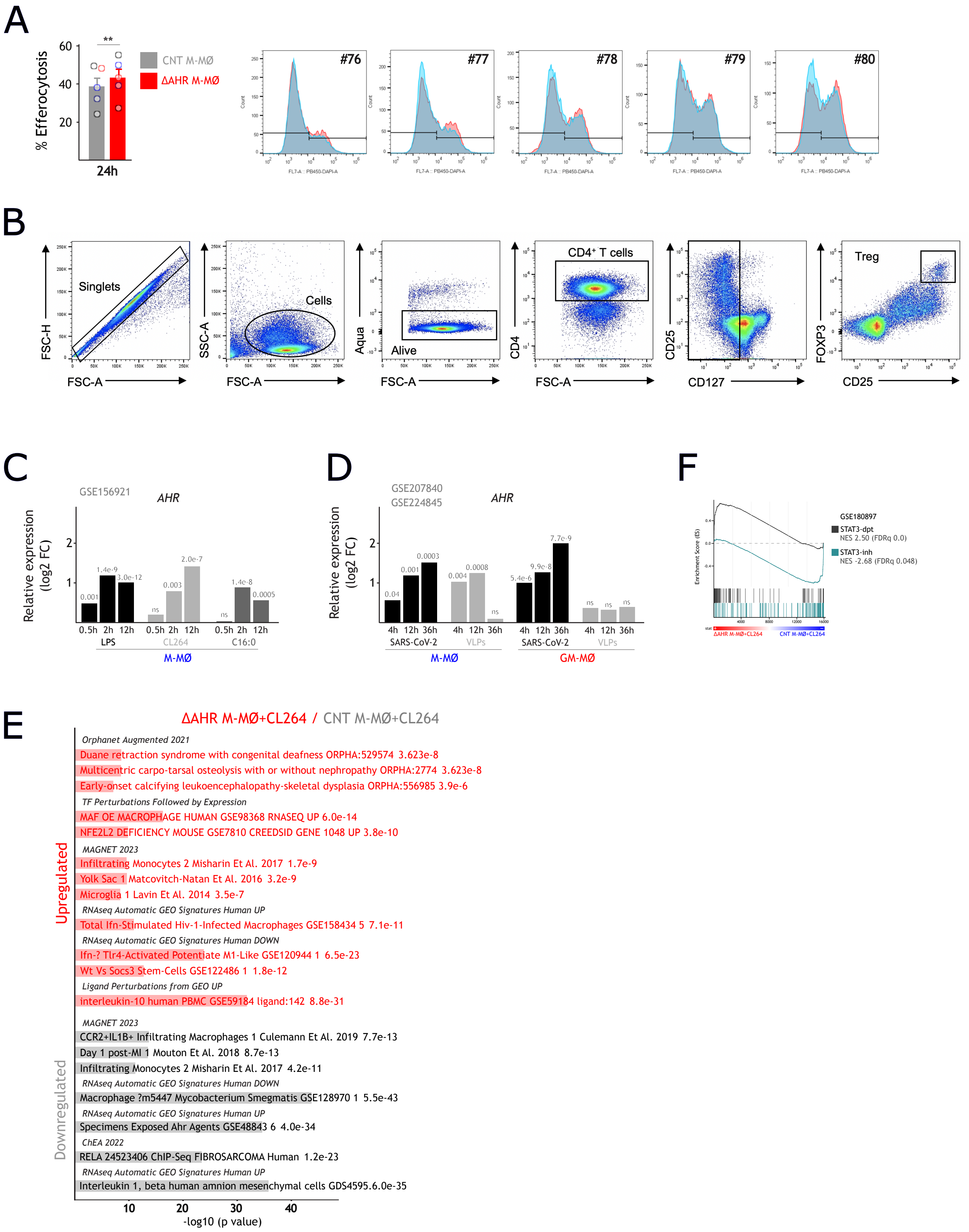


Supplementary Figure 4

**Supplementary Figure 4.- Functional consequences of AhR silencing. A.** Quantification and flow cytometry profiles (CellTrace Violet emission) of the efferocytosis capacity of ∆AHR-M-MØ and CNT M-MØ, determined 24 h after siRNA transfection (n=5; mean ± SEM; paired Student´s t-test: **, p<0.01). **B**. Gating strategy of $CD4^{+}$ $CD127^{-}$ $CD25^{high}$ $FOXP3^{high}$ Tregs induced by allogeneic CNT M-MØ or ΔAHR M-MØ after 5 days of coculture. **C.** *AHR* mRNA expression in M-MØ stimulated with LPS, CL264 or palmitate (C16:0) at the indicated time points (from GSE156921) [5141]. **D**. *AHR* mRNA expression in M-MØ or GM-MØ exposed to SARS-CoV-2 or Virus-like particles (VLPs) at the indicated time points (from GSE207840 and GSE224845) [21]. In **C/D**, statistical significance of the comparison to untreated cells is indicated (ns, not significant). **E**. Enrichr [61] gene ontology analysis of the genes differentially expressed between CL264-treated ∆AHR-M-MØ and CL264-treated CNT M-MØ. **F**. GSEA profiles of STAT3-regulated genes (GSE180897) on the ranked comparison of CL264-treated ∆AHR-M-MØ vs. CL264-treated CNT M-MØ transcriptome. NES and FDRq values are indicated.

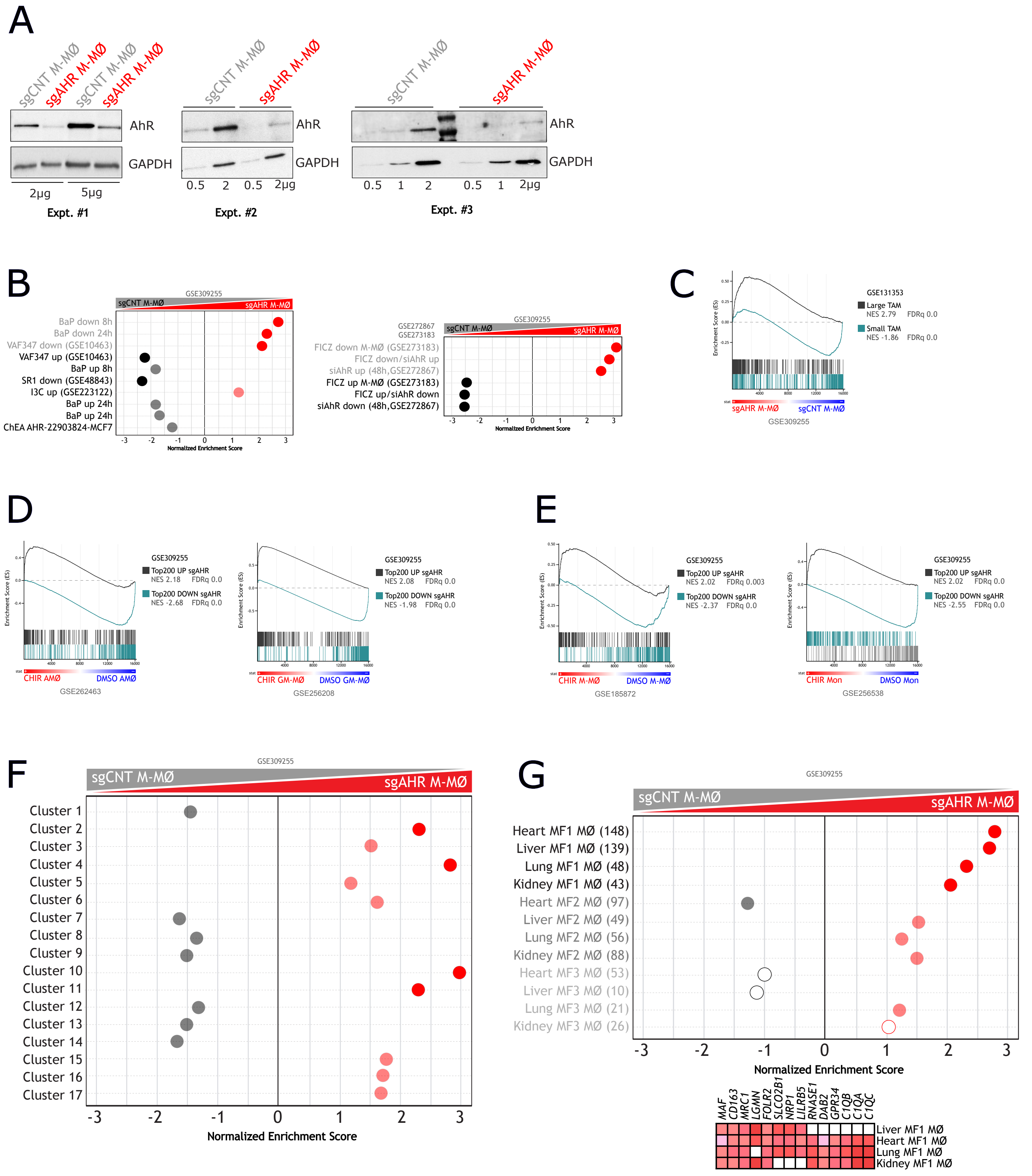


Supplementary Figure 5

**Supplementary Figure 5.- AhR ablation in monocytes skews macrophage differentiation. A.** Western blot of AhR on the indicated amount of cell lysate from three independent preparations of sgCNT M-MØ and sgAHR M-MØ (n=3). **B**. GSEA summary of canonical AhR-regulated gene sets [from [3233343536]] (*Left panel*), and defined in this manuscript (*Right panel*), on the ranked comparison of the sgAHR-M-MØ vs. sgCNT M-MØ transcriptome. NES and FDRq values are indicated (solid colors, FDRq=0; light colors, FDRq<0.25). **C**. GSEA profiles of the gene sets that define "Large TAM" and "Small TAM" from colorectal liver metastasis on the ranked comparison of the sgAHR-M-MØ vs. sgCNT M-MØ transcriptome. NES and FDRq values are indicated. **D/E**. GSEA profiles of the Top200 genes upregulated (**D**) or downregulated (**E**) in sgAHR-M-MØ on the ranked comparison of the transcriptomes of CHIR-AMØ vs. DMSO-AMØ, CHIR-GM-MØ vs. GM-MØ, CHIR-M-MØ vs. M-MØ, and CHIR-Mon vs. DMSO-Mon. NES and FDRq values are indicated. **F**. Summary of GSEA of the gene sets that define the MoMac-VERSE clusters [5] on the ranked comparison of the sgAHR-M-MØ vs. sgCNT M-MØ transcriptome. NES and FDRq values are indicated (solid colors, FDRq=0; light colors, FDRq<0.25). **G**. GSEA (*Left panel*) and leading-edge analysis (*Right panel*) of the gene sets that define the three subsets of tissue-resident macrophages [as defined in [55]] on the ranked comparison of the sgAHR-M-MØ vs. sgCNT M-MØ transcriptome. NES and FDRq values are indicated (solid colors, FDRq=0; light colors, FDRq<0.25; Not significant, empty circles).

Supplementary Table S2.
Transcriptional datasets used in the present study

| Reference | Sparfel et al. Toxicol Sci 114:247, 2010 | | | | Lawrence et al. Blood 112:1158, 2008 | | Pabst et al. Blood 127:2018, 2016 | Schmidt et al. Front Immunol 14, 2023 | Lo et al. Toxicol Sci 130:349, 2012 | Grant et al Nature 590:635, 2021 | | | | | | Wendisch et al Cell 184:6243, 2021 | |
|---|---|---|---|---|---|---|---|---|---|---|---|---|---|---|---|---|---|
| Repository | | | | | GSE10463 | | GSE48843 | GSE223122 | | GSE155249 | | | | | | EGAS00001005634 | |
| Name | BaP down 8h | BaP down 24h | BaP up 8h | BaP up 24h | VAF347 up | VAF347 down | SR1 down | I3C up | ChEA AHR-22903824-MCF7 | MoAM1 | MoAM2 | MoAM3 | MoAM4 | TRAM1 | TRAM2 | Monocytes | Monocyte/MØ |
| Genes | ENAH | STK39 | A4GALT | SERPINB2 | THBS1 | CHI3L2 | CYP1A1 | SERPINB2 | CYP1A1 | ACSL1 | ACSL1 | ABCA1 | AGTRAP | ABCG1 | ABCG1 | AC245128.3 | ACSL4 |
| | UBE2C | MT1E | AHRR | CXCR5 | PLAT | GOLGA7B | CYP1B1 | MMP27 | CYP1A2 | ANKRD22 | ADAP2 | ABHD12 | AIF1 | ABHD5 | ACOT2 | ACTB | ACTR2 |
| | FAM64A | SDCCAG10 | AKR1C1 | IL1B | IL19 | RPL36A | AHRR | PTGS1 | CYP1B1 | ANXA5 | AIF1 | ACP2 | ANXA2 | AC026369.3 | ACP5 | ACTG1 | AIF1 |
| | LIMA1 | EXO1 | ARMC9 | PLK2 | AHRR | SERPINF1 | IL9R | AHRR | ALDH1A3 | APOBEC3A | ANKRD22 | ADAP2 | ANXA5 | ACOT2 | ACSL1 | ADAM19 | ALCAM |
| | NDC80 | TPK1 | ASB2 | IL8 | RASAL1 | MRPL17 | ITGA1 | MMP8 | TIPARP | AQP9 | ANXA5 | AIF1 | AP2S1 | ACP5 | ACTN1 | AIF1 | ANKRD22 |
| | EXO1 | TYW1 | ATP1B2 | ATP1B2 | SEMA6B | COX17 | EPGN | SPINT1 | AHRR | ATF3 | APOBEC3A | ANXA2 | APLP2 | ADAMTSL4 | ADTRP | AL034397.3 | APP |
| | TOP2A | NUBPL | CCL1 | RASAL1 | XYLT1 | DCTPP1 | AMZ1 | TCF19 | SLC7A5 | ATF5 | ATF5 | ANXA5 | APOBEC3A | AGPAT2 | AGRP | APOBEC3A | APRT |
| | LRRC25 | PRKCB | CDK5RAP2 | EREG | IL24 | CHCHD10 | F2RL3 | NQO1 | WDR63 | ATOX1 | ATG3 | AP1B1 | AQP9 | AGRP | AKR1B1 | ARHGDIB | AQP9 |
| | FAM19A3 | AK5 | CDKN1A | GADD45A | FRY | SERPINI1 | EPB41L3 | EMP1 | SECTM1 | ATP2B1 | ATOX1 | AP2A2 | ARHGAP18 | AKR1B1 | ALAS1 | ASGR2 | ARPC5 |
| | P2RY8 | SAMD4A | CFP | AHRR | CYP1B1 | ZNF786 | GPR68 | CD28 | DLX2 | ATP6V1B2 | BAX | AP2M1 | ARPC3 | AKR1C3 | ALDH1A1 | BACH1 | ATP6V0D1 |
| | CCNA2 | MYO9A | CST7 | GPR35 | LRG1 | NDUFA3 | CCL24 | CLIC5 | C21orf29 | ATP6V1F | BCL2A1 | AP2S1 | ASGR1 | ALDH1A1 | ALDH2 | C3AR1 | ATP6V1G1 |
| | RGS18 | GINS2 | CXCL1 | WNT5A | ATP1B2 | ADRB2 | DMTN | P2RY13 | BMF | ATP6V1G1 | BLVRA | APLP2 | ATOX1 | ALDH2 | ALOX5 | CCDC69 | BASP1 |
| | E2F2 | HIST1H1B | CXCL3 | SNAI1 | FCAR | IFIT2 | SEMA6B | ACPP | LRRC15 | BACH1 | BLVRB | APOC1 | ATP5F1E | ALDOA | ALOX5AP | CCR2 | CALM1 |
| | KIF20A | PNPT1 | CXCR5 | DPYSL3 | LPAR6 | GP1BA | TGM5 | TMEM37 | RND1 | BCL2A1 | BRI3 | APOE | ATP6V0B | ALOX5 | ANXA2 | CD14 | CCL2 |
| | FAM105A | ATPBD4 | CYP1B1 | F2RL2 | TMEM45B | CCL26 | NPTX1 | LPAR6 | VIPR1 | BCL3 | BST2 | ARHGAP18 | ATP6V0C | ALOX5AP | ANXA5 | CD55 | CCR2 |
| | DTL | TULP4 | DPYSL3 | KRT23 | TIPARP | FCGR1B | EREG | CTLA2B | EDC3 | BRI3 | C1orf162 | ASAH1 | ATP6V0D1 | AMIGO2 | APIP | CD93 | CCR5 |
| | MYC | E2F2 | ENC1 | VASN | GPR68 | NDUFA6-AS1 | NCF1B | GM12185 | MCOLN2 | C1orf122 | C3AR1 | ATOX1 | ATP6V1F | APOC1 | APOC1 | CD300E | CD44 |
| | SLA | MT1A | EREG | AKR1C1 | CSRP2 | C4orf48 | CXCL8 | MTFR2 | ST3GAL1 | C3AR1 | C15orf48 | ATP5F1E | BCL2A1 | APOE | APOE | CDA | CD93 |
| | NLRP12 | MT1H | FCAR | NIACR2 | RCBTB2 | MARCO | IL1A | ANGPTL2 | LMCD1 | C5AR1 | CALHM6 | ATP5PD | BLVRB | ASAH1 | ARRDC4 | CEBPD | CD300E |
| | ZC3HAV1L | DOCK10 | FUCA1 | FDXR | LACC1 | OCEL1 | CNR2 | GM6545 | PYGL | C15orf48 | CALM3 | ATP6V0B | BRI3 | ATP1B1 | ASAH1 | CFP | CEP170 |
| | TRIB3 | GMDS | GALNTL4 | SEMA6B | BHLHE41 | PTRHD1 | THBS1 | DNAH2 | RUNX2 | CCL2 | CASP4 | ATP6V1F | C1orf162 | BCAP31 | ATP1B1 | CKAP4 | CLEC5A |
| | CENPA | BRCA1 | GCLM | CDK5RAP2 | LOC100996412 | SLIRP | IL1B | FAM84B | LMO2 | CCL3 | CBR1 | BAX | C4orf48 | BHLHE41 | ATP6AP1 | CLU | CLEC10A |
| | MT1L | MYC | GFPT2 | CFP | FCMR | PRIM1 | SERPINB2 | GGTA1 | SIPA1L2 | CCL3L1 | CCL2 | BLVRB | C15orf48 | C1orf162 | ATP6V0C | CORO1A | CLU |
| | MT1G | UBE2E2 | GNG11 | RARRES2 | SLC5A3 | RPL22L1 | NCF1C | VCAN | | CCL4 | CCL7 | BMP2K | CALHM6 | C1QA | AXL | COTL1 | CORO1A |
| | MT1H | PTK2 | GPR183 | IL1A | ARL4C | AMPD2 | NCF1 | RHOV | | CCL4L2 | CCL8 | BRI3 | CALM1 | C1QB | B2M | CRISPLD2 | CYBB |
| | UHRF1 | FOXP1 | GPR35 | EID3 | TRERF1 | CCL27 | ABI3 | GSDME | | CCL7 | CCR1 | C1orf162 | CAPG | C1QC | BCAP31 | CSF3R | DDX5 |
| | MT1B | PRIM2 | GPR64 | CST7 | PITPNC1 | LINC00869 | LOC100288175 | GM20186 | | CCL8 | CD14 | C1QA | CCL2 | C9orf16 | BCL2A1 | DSE | DDX21 |
| | MT2A | THRB | GPR68 | ACTA2 | CCL1 | ALDH3A1 | RMDN2 | GSTA3 | | CCR1 | CD63 | C1QB | CCR1 | CAT | BLVRA | DUSP6 | DNAJC8 |
| | MT1A | CENPP | IER3 | CCL1 | RUNX2 | CKS1B | TIPARP | P2RY14 | | CCRL2 | CD84 | C1QC | CD14 | CCDC88A | BRI3 | EFHD2 | DPYD |
| | MT1X | EXOC4 | IL1A | SLC16A6 | THBD | TOR3A | MGAT3 | LY6A | | CD14 | CD163 | CAPG | CD44 | CD9 | C1QA | EGR1 | DSE |
| | MT1E | KCNMA1 | IL1B | CABYR | EHF | TSTD1 | MMP25 | ABCB4 | | CD44 | CD300A | CCL2 | CD63 | CD52 | C1QB | EIF1 | ELK3 |
| | MT1F | MT1L | ITGB8 | GDF15 | ADA | ACY1 | UNC5B | SERPINB8 | | CD48 | CD300E | CCL18 | CD68 | CD68 | C1QC | ELK3 | EMP1 |
| | TMEM37 | UBE2CBP | LEPREL1 | DDB2 | PKIB | DTX4 | MMP25-AS1 | C130050O18RIK | | CD63 | CDKN1A | CCR1 | CD93 | CD74 | C3AR1 | FAU | EMP3 |
| | CDCA7 | CEP55 | LRG1 | GNG11 | SLC9A9 | BATF3 | LOC105378948 | E2F8 | | CD83 | CEBPD | CD14 | CD163 | CD81 | C5AR1 | FCN1 | FAM49B |
| | MT1M | CDYL2 | NCF1 | FAS | FAM20C | SNORA24 | RUNX3 | PGAP1 | | CD300E | CFL1 | CD36 | CD300E | CD151 | CAPG | FGL2 | FCGR2B |
| | [illegible] | [illegible] | [illegible] | [illegible] | [illegible] | [illegible] | [illegible] | [illegible] | | [illegible] | [illegible] | [illegible] | [illegible] | [illegible] | [illegible] | [illegible] | [illegible] |

[illegible]

| |
|---|
| C1orf63 |
| THAP2 |
| SPATA18 |
| TNFRSF10C |
| ZNF79 |
| RORA |
| LEPREL1 |
| ZNF425 |
| ADORA2B |
| PTP4A1 |
| RIT1 |
| UBIAD1 |
| GPR175 |
| RUNX3 |
| C19orf40 |
| MFSD4 |
| BCLAF1 |
| PIM2 |
| PCNA |
| ST6GALNAC2 |
| TRAF1 |
| PTGR1 |
| UBR5 |
| LRRC8E |
| PSTPIP2 |
| HIST1H2BD |
| BEX1 |
| NAMPT |
| ARRDC4 |
| SNX16 |
| CDC37L1 |
| GRPEL2 |
| DNAJB9 |
| SDCBP2 |
| ANKRA2 |
| MYLIP |
| PPP1R15B |
| TMEM22 |
| WWOX |
| TTRAP |
| ANGPTL6 |
| GPC1 |
| FAM3C |
| CCRK |
| MXD1 |
| ERCC6 |
| RRM2B |
| PNPO |
| TSKU |
| EIF5 |
| HIST1H1C |
| OTUD1 |
| RBBP6 |
| IDO1 |
| LYSMD1 |
| RRAS2 |
| PERP |
| MSC |
| DNAJB4 |
| HIST1H2BO |
| C17orf100 |
| PPIF |
| C1RL |
| NFYA |
| FLJ43855 |
| RGAG4 |
| C21orf121 |
| KIAA0323 |
| SLFN5 |
| RGS12 |
| SYNC |
| TMEM217 |
| DCP1B |
| TNFAIP3 |
| ADAMDEC1 |
| C3orf71 |

[illegible]

| | | | |
|---|---|---|---|
| SPINT2 | ACE | KYNU | FAR2 |
| FOLR2 | ENC1 | SCIN | ZNF268 |
| EPOR | CRYBB1 | UBE2Z | SH3RF3 |
| PINK1 | MXD1 | NAMPT | RGS3 |
| SPINT1 | HRH1 | APOBEC3G | CBWD3 |
| NHSL2 | ANTXR1 | G3BP1 | CLEC5A |
| FOXO1 | RHOBTB1 | AKT1S1 | AKR1C1 |
| TDP1 | OLFML2B | AKAP2 | CBS |
| NAV1 | CXCL12 | TMEM138 | LILRA1 |
| ACBD5 | F2RL3 | ATP13A2 | GPAT3 |
| ABCA7 | LRRC4 | MTHFD2 | MAP2K6 |
| PTGS1 | MERTK | B4GALT5 | EDNRB |
| ATG16L2 | SEMA4C | TUFT1 | TMEM158 |
| SUSD1 | ACER2 | DCANP1 | CD72 |
| BACE1 | ZNF704 | IFIH1 | SLC12A8 |
| DGKD | NBL1 | CGAS | NCAM1 |
| CEP85L | THBS1 | TNFRSF18 | FNTB |
| CORO2A | TMEM26 | CDKN1A | PIM1 |
| JMJD7-PLA2G4B | TMIGD3 | PIWIL4 | CLEC12A |
| CIDEB | HS3ST1 | CCL7 | APOBEC3B |
| ZNF652 | COLGALT2 | USP18 | SULT1A3 |
| GPRIN3 | LGMN | WTAP | MYBPH |
| KANK2 | AATK | KANK1 | TRPV4 |
| PGM1 | LRP5 | NFKB2 | CD1D |
| RMND5A | SLC22A23 | | |
| FES | RBP1 | | |
| CHID1 | GFRA2 | | |
| VSIR | ITGA9 | | |
| DENND6B | CD28 | | |
| NOMO2 | SERPINB2 | | |

[illegible]

| |
|---|
| TUBB4B |
| TUBB6 |
| TUBG1 |
| TYMS |
| UBE2C |
| UBE2S |
| UBE2T |
| UHRF1 |
| USP1 |
| VDAC3 |
| VPS29 |
| VRK1 |
| WDHD1 |
| WDR34 |
| WDR62 |
| WDR76 |
| WEE1 |
| XRCC2 |
| YWHAH |
| ZDHHC12 |
| ZGRF1 |
| ZNF93 |
| ZWILCH |
| ZWINT |

| |
|---|
| KRT79 |
| ARHGEF28 |
| FN1 |
| INHBA |
| APOL4 |
| AQP3 |
| STAC |

| |
|---|
| MTM1 |
| TENT2 |
| TNFSF8 |
| SLC31A2 |
| LGALS9 |
| IFIT5 |
| PLEKHO2 |
| PTPRC |
| CCND3 |
| SNX6 |
| ADAMTSL4 |
| PELI1 |
| SNX20 |
| RERE |
| VSIG4 |
| TTC38 |
| PTK2B |
| MSL3 |
| CBR1 |
| PRELID1 |
| SLAMF7 |
| WDR1 |
| EMP1 |
| PTPRJ |
| ITPRID2 |
| METRNL |
| ABI1 |
| NECTIN2 |
| STAT6 |
| ELF1 |
| GIMAP1 |
| CLIC4 |
| TPM3 |
| MYL12A |
| PCGF5 |
| IGSF6 |
| DCK |
| C3AR1 |
| STING1 |
| TREX1 |
| SNX2 |
| ZFP36L2 |
| ANXA4 |
| STAT2 |
| HLA-A |
| TMEM63A |
| B4GALT5 |
| CYRIB |
| NFE2L1 |
| XAF1 |
| IER2 |
| RGL1 |
| IFI44L |
| SGMS1 |
| RNASET2 |
| WIPF1 |
| EXT2 |
| ST8SIA4 |
| TMSB4X |
| PIK3CD |
| OGFR |
| TNFSF13 |
| TENT4A |
| B4GALT1 |
| BTG1 |
| CMTR1 |
| IFI44 |
| FCGR2A |
| CORO1A |
| MPP1 |
| OPTN |
| DYNC1LI2 |
| ERP44 |
| MAX |
| ADCY7 |
| HLA-B |
| HDAC6 |
| ADAR |
| COMT |
| LGMN |
| HSPA6 |
| SP100 |
| ARHGAP25 |
| HPS5 |
| DNAJA1 |
| IRF2BPL |
| PDIA4 |
| SAP18 |
| MCFD2 |
| FNBP1 |
| ZFP36 |
| PATL1 |
| TRIM38 |
| TMEM123 |
| SYNGR2 |
| INPP5D |
| LPCAT1 |
| TRIM26 |
| FYB1 |
| TRPM2 |
| FKBP15 |
| HNMT |
| ATP13A1 |
| FLI1 |
| SNN |
| CASP4 |
| RNF114 |
| FRMD4A |
| HCK |
| TESK1 |
| PRR13 |
| IL10RA |
| HIF1A |
| NADK |
| TRPC4AP |
| SIGLEC1 |
| RB1 |
| GPR137B |
| CTSL |
| HCLS1 |
| KPNB1 |
| TCN2 |
| HAVCR2 |
| CSF3R |
| TNFSF13B |
| RNASE1 |
| STAT3 |
| TFEC |
| RAC1 |
| CCR1 |
| ITM2B |
| GRK2 |
| SASH3 |
| C1QA |
| C2 |
| MOB1A |
| MSN |
| CSF1R |
| GLUL |
| GNAI2 |
| B2M |

| |
|---|
| KIF18B |
| CCR2 |
| MMAB |
| LIPA |
| PAQR5 |
| NES |
| CHIT1 |
| DUSP2 |
| FADS1 |
| LSP1 |
| SGMS2 |
| TFAP2A |
| ARAP3 |
| RMI2 |
| USP2 |
| GPC4 |
| NFE2L3 |
| QPCT |
| TNFRSF9 |
| GPR55 |
| KNL1 |
| HMGCR |
| SIGLEC16 |
| PRC1 |
| PFKP |
| ENTHD1 |
| CKAP4 |
| VIT |
| KCNN4 |
| ADRA2B |
| PHLDA3 |
| CXCL1 |
| SHCBP1 |
| CCND3 |
| DIRAS2 |
| FOXM1 |
| NEURL1B |
| PIF1 |
| AKR1C2 |
| CCL22 |
| FAM81A |
| CCL1 |
| ATF3 |
| GINS2 |
| PNPLA3 |
| U2AF1L5 |
| DCSTAMP |
| LPCAT2 |
| C20orf197 |
| PNP |
| HELLS |
| FASN |
| CCL8 |
| SSTR2 |
| SPTBN1 |
| ACOT7 |
| NME1 |
| IQGAP3 |
| CRABP2 |
| ALCAM |
| OLR1 |
| ADAM12 |
| ITGB7 |
| CDKN3 |
| CDCA3 |
| CCND1 |
| POLQ |
| TM4SF19 |
| CLEC6A |
| CYP51A1 |
| MEIS3 |
| IL1B |
| ZNF367 |
| TCTEX1D1 |
| SH3RF3 |
| RAC2 |
| PKIB |
| CENPF |
| DHCR7 |
| SQLE |
| PROK2 |
| SCD |
| SPC24 |
| HPGDS |
| SLC30A3 |
| NUF2 |
| CIT |
| ASPHD1 |
| ZBTB7C |
| CNIH3 |
| SGO1 |
| TPRG1 |
| CLDN1 |
| MGAM |
| MATK |
| PIMREG |
| TNFAIP8L3 |
| HSD11B1 |
| TGM2 |
| TTC39B |
| SPAG5 |
| RAD51 |
| OCSTAMP |
| TROAP |
| NCS1 |
| ST20 |
| ALOX5AP |
| CENPM |
| CREB5 |
| NEDD9 |
| PALD1 |
| BUB1 |
| MSMO1 |
| PCDHGC3 |
| PTX3 |
| SAPCD2 |
| INHBA |
| PHGDH |
| CDCA5 |
| CCNA2 |
| ESPL1 |
| KIF23 |
| C6orf223 |
| CD300E |
| KNDC1 |
| CDCA7 |
| PLK1 |
| MGST1 |
| MELK |
| CENPK |
| ALDH1A2 |
| TTK |
| CCL4L1 |
| GALNT12 |
| RGCC |
| MAMLD1 |
| BCL2A1 |
| CCNB1 |
| GPAT3 |
| CDCA2 |
| RAD54L |
| AKAP2 |
| DEPDC1B |
| HJURP |
| ADGRE3 |
| PROCR |
| CA12 |
| ANLN |
| CD52 |
| KIF4A |
| UBE2C |
| KIF2C |
| CDC6 |
| APOBEC3B |
| HMMR |
| IGSF21 |
| UHRF1 |
| CENPA |
| TRIP13 |
| E2F7 |
| E2F2 |
| CEP55 |
| SCG5 |
| PLEK2 |
| KIFC1 |
| SLC28A3 |
| TICRR |
| CDC25C |
| FADS2 |
| E2F1 |
| RIMBP3C |
| LRP8 |
| NCAPH |
| PCSK6 |
| KIF14 |
| CCNB2 |
| TPX2 |
| CCL2 |
| IL1RN |
| CDC45 |
| SLC2A5 |
| NRGN |
| SLAMF9 |
| CYP19A1 |
| DIAPH3 |
| TOP2A |
| CKAP2L |
| DEPDC1 |
| LPL |
| CD1E |
| NEIL3 |
| CSF1 |
| MYOF |
| ASPM |
| STAC |
| BUB1B |
| DHCR24 |
| SLIT3 |
| EXO1 |
| GTSE1 |
| IGSF10 |
| NDC80 |
| CD1C |
| SKA3 |
| CDK1 |
| CDC20 |
| ESCO2 |
| CHI3L1 |
| KIF15 |
| DLGAP5 |
| SPN |
| FBP1 |
| AURKB |
| FAM83D |
| CCNE2 |
| RTN1 |

| |
|---|
| MYLIP |
| MEF2C |
| ITGAL |
| SPATA41 |
| CEBPG |
| LOC644656 |
| SRSF5 |
| TGIF1 |
| NCOA5 |
| TIMM44 |
| FOSB |
| C16orf72 |
| MDM4 |
| NAB1 |
| TOMM5 |
| YPEL5 |
| COQ7 |
| LOC284454 |
| NFKBIZ |
| NOP56 |
| TPT1 |
| FAM219B |
| EFR3A |
| SENP8 |
| PIK3CD-AS1 |
| ASAH1 |
| CEBPD |
| HMG20A |
| PSMA1 |
| CALM1 |
| AFF1 |
| INCENP |
| NRDC |
| ASB16-AS1 |
| PTPRE |
| LOC105376805 |
| LAPTM5 |
| RSL24D1 |
| SLC36A4 |
| PFN1 |
| PYCR2 |
| PFKFB3 |
| BNIP2 |
| MORF4L2-AS1 |
| YWHAZ |
| DHPS |
| CHMP4A |
| FAM107B |
| OAZ1 |
| PCBP2 |
| LACC1 |
| TIPARP |
| PER1 |
| FLJ10038 |
| PI4KA |
| FSCN1 |
| KCTD12 |
| PRMT3 |
| SIRT4 |
| CTNNB1 |
| PSMB5 |
| RELT |
| NAGK |
| ZFP36L2 |
| NORAD |
| SIRT1 |
| NR4A2 |
| CAST |
| CEBPA |
| IRS2 |
| MKNK2 |
| HNRNPA0 |
| IARS |
| SLC19A2 |
| ZNF302 |
| STEAP4 |
| NOL11 |
| NABP1 |
| NIT1 |
| TPM3 |
| MED28 |
| DDIT4 |
| EPAS1 |
| SIPA1L1 |
| SLFN11 |
| TCN1 |
| C15orf39 |
| ADAM9 |
| TNFRSF1A |
| CD55 |
| STX11 |
| RNU4-1 |
| MALT1 |
| PLSCR1 |
| ICAM1 |
| PLAUR |
| ARIH2 |
| CIRBP |
| KCTD5 |
| SMURF2 |
| THEMIS2 |
| PPIB |
| MANBA |
| ID2 |
| TAL1 |
| KIAA1033 |
| LOC101928132 |
| DMXL1 |
| DDX5 |
| NUFIP2 |
| NAPA |
| P2RY12 |
| RALY |
| DDX3X |
| GPX4 |
| SNHG25 |
| DDRGK1 |
| FADS1 |
| HAL |
| HNRNPA2B1 |
| RAB31 |
| RNU11 |
| TNPO1 |
| SLC9B1 |
| RPLP0 |
| USP36 |
| ZCWPW2 |
| HBEGF |
| MEFV |
| SNHG7 |
| C20orf197 |
| DYNLL2 |
| VAMP1 |
| EHF |
| TIMP2 |
| RIPK1 |
| SENP7 |
| WDR74 |
| TFRC |
| COQ5 |
| LOC100289495 |
| RPN1 |
| ZFAND5 |
| DNAJB9 |
| TRUB1 |
| GRB2 |
| FAM227B |
| FFAR4 |
| FUS |
| LOC647323 |
| SRSF7 |
| ZCWPW1 |
| ZNF548 |
| ACTR2 |
| CAB39 |
| DBP |
| RANBP9 |
| ARL4C |
| DNAJC25 |
| LINC00968 |
| ARL6IP1 |
| C5AR1 |
| HN1 |
| KLHL24 |
| SDCBP |
| TTYH3 |
| SEPP1 |
| CYSTM1 |
| ELP3 |
| CARHSP1 |
| DNAJA1 |
| EHD1 |
| TGOLN2 |
| AFF4 |
| KLF7 |
| MTRR |
| ZNF331 |
| DNAJC27 |
| GRAMD3 |
| PABPC1 |
| SEC24B |
| XRN1 |
| AOAH |
| CHMP1B |
| EIF4A2 |
| EXOSC9 |
| RNU5B-1 |
| RPS27 |
| TMEM52B |
| HSP90AA1 |
| LINC00467 |
| MYL6 |
| NARF |
| TRAPPC6A |
| GPX1 |
| IL10RB-AS1 |
| MTO1 |
| OSER1 |
| SETDB2-PHF11 |
| ATF4 |
| CFL1 |
| CHMP2A |
| CITED2 |
| CPSF6 |
| HEXB |
| RCOR1 |
| FOSL2 |
| RHOB |
| SPTAN1 |
| NFIA |
| GALNT11 |
| LOC101929741 |
| LOC285074 |
| NT5C2 |
| NXF1 |
| SRGN |

| | |
|---|---|
| ZFHX3 | KCNAB2 |
| NR1H3 | MIER2 |
| RGCC | TMEM176A |
| SEPTIN11 | FAM135A |
| TUBA1B | RAB11FIP1 |
| SORBS3 | TNFAIP1 |
| ADGRE5 | HTT |
| C15orf48 | ALDH2 |
| CD300A | FLYWCH1 |
| ITGA3 | NUDT4 |
| ACSL1 | MIGA2 |
| CKLF | APOBR |
| RAPH1 | IL10RB |
| FGD6 | NLRX1 |
| MCM3 | ADCY9 |
| CXCL2 | SLC43A2 |
| TCF4 | PDE4DIP |
| PGM1 | FNDC3A |
| DENND2D | CRIM1 |
| ABRACL | SRGN |
| ARMT1 | APBA1 |
| FGFR1 | MPPE1 |
| LDLRAD4 | PDCD6 |
| DDX60L | GNA12 |
| MAML2 | SKI |
| CXCR4 | SELPLG |
| SLC39A8 | RAB27A |
| ADAP2 | SERINC5 |
| FPR1 | FCGR2B |
| LCP2 | DGKQ |
| RARRES1 | VSIR |
| IER2 | SARAF |
| ADARB1 | ZBTB4 |
| SIGLEC14 | ARHGAP22 |
| PIK3AP1 | CAMK1 |
| DUSP2 | ANTXR2 |
| RASGRP3 | MAN2A2 |
| DUSP1 | POLR2E |
| RTN1 | NIBAN2 |
| EMB | ATP13A3 |
| PAPSS2 | IRF1 |
| FASN | INPP4A |
| PNP | IRS2 |
| KLF3 | LIMD1 |
| FCMR | SLC12A7 |
| BMP2K | ITGB5 |
| GIMAP6 | IDH1 |
| C6orf62 | WDR45B |
| MYO19 | PPM1F |
| ST3GAL6 | SDHA |
| CD276 | ARHGAP45 |
| RPP25 | SLC3A2 |
| TSC22D1 | RABAC1 |
| LPCAT1 | ATG2A |
| TBC1D4 | PTTG1IP |
| HPRT1 | MPEG1 |
| NME1 | TOM1 |
| SNX24 | STAT1 |
| CISH | CLN8 |
| NCOA7 | CYP2S1 |
| PANX1 | FOXN2 |
| CA2 | YWHAZ |
| PEAK1 | RHBDF2 |
| TMEM37 | GPATCH2L |
| FADS1 | RGS10 |
| EGFL7 | LIMS1 |
| SSBP2 | ALAS1 |
| RAP2B | FPR3 |
| SUCNR1 | UNC13D |
| TRAF3IP3 | JPT1 |
| HSPB1 | PARL |
| PRKCB | CD93 |
| HRH2 | GABARAPL1 |
| VAV3 | SLC31A2 |
| BCL6 | AP5B1 |
| CKAP4 | NAIP |
| ITGAX | BCL3 |
| AFAP1L1 | KIAA1191 |
| RNF145 | COLGALT1 |
| NFKBIZ | RELT |
| PLCXD1 | RAD23A |
| SLC8A1 | DPP9 |
| CCDC170 | WARS1 |
| GLIPR2 | ACSS2 |
| HECTD2 | HBP1 |
| DHFR | CTSO |
| BHLHE40 | CLCN7 |
| C3AR1 | SLC2A9 |
| CD82 | SLC2A3 |
| CYP51A1 | MEGF9 |
| IGSF6 | PGS1 |
| ACRBP | TMEM176B |
| LOC102724250 | TMEM50A |
| CBLB | CALR |
| C1QB | GRINA |
| MSR1 | DUSP3 |
| SPOCD1 | SLC7A8 |
| SCMH1 | SMARCD2 |
| NPIPB3 | MAPKAPK2 |
| DEF6 | ANKH |
| FBP1 | TMEM170B |
| LGALS9 | SPI1 |
| C1orf127 | GPX1 |
| MREG | S100A4 |
| TM4SF19 | PPP2R5D |
| NCKAP1 | KCTD10 |
| ZNF467 | MYO1F |
| CNIH3 | RIPK1 |
| JAKMIP2 | CAT |
| PDE4B | CAPZB |
| CPM | MGRN1 |
| RHOBTB1 | USP4 |
| MS4A4E | PRDM1 |
| FOXP1 | SLC36A1 |
| ZHX2 | SERPINB1 |
| CDC42EP3 | RASSF5 |
| STARD8 | DNASE2 |
| AP1S2 | HPS3 |
| GAS2L3 | AP1B1 |
| SMCO4 | CUX1 |
| LXN | SLC26A11 |
| TUBB | MAP1LC3B |
| LFNG | S100A6 |
| DNAJC9 | PFKFB3 |
| NRIP3 | ATP8B4 |
| NUP210 | EDEM2 |
| SNCA | PPP2R5C |
| MAP7 | CHPF2 |
| NPIPB5 | MAPK1 |
| MARCO | IL6R |
| PLK3 | MBNL1 |
| MEF2C | VPS4B |
| KPNA2 | RRM2B |
| CLEC2D | MTMR3 |
| B3GNT5 | CYTH1 |
| ABCC5 | NCOA3 |
| SAT1 | FAM78A |
| SAMD4A | IER3 |
| HPGD | RCOR1 |
| EGR1 | DNM2 |
| OSBPL3 | SEC62 |
| RGS2 | HSPA6 |
| FOXRED2 | SUPT5H |
| PDLIM2 | IL4R |
| FAM20A | PSD4 |
| TLE3 | RIN2 |
| HAVCR2 | EMILIN1 |
| HLA-DOA | CCNL1 |
| PLCL1 | LMAN2 |
| GPAT3 | CD86 |
| ALCAM | SIRT2 |
| PCSK6 | MCMBP |
| ITPKB | GPX4 |
| MTSS1 | SEC16A |
| HFE | PGD |
| EPAS1 | ZNF706 |
| TIMP3 | GPNMB |
| ID2 | PRELID1 |
| PLEK | SH3GL1 |
| SLC41A2 | DGCR2 |
| HAGHL | KHNYN |
| KCNN4 | AP2A2 |
| CD83 | GNPTG |
| MAN1C1 | GRAMD4 |
| CDK6 | VPS26B |
| PPFIBP1 | PLEKHO1 |
| CACNA2D3 | RIN3 |
| ARAP2 | ABHD4 |
| CADM1 | SLC37A2 |
| DYSF | IL13RA1 |
| EDA | TYK2 |
| PIK3R6 | RAPGEF1 |
| ITSN1 | SBF1 |
| SGK1 | ENG |
| ANTXR1 | SYNGR2 |
| ARHGAP24 | IRF2 |
| ZDHHC14 | RCSD1 |
| NBEAL2 | CD180 |
| POGLUT3 | DAZAP2 |
| VAMP5 | RNASE6 |
| GRAMD1B | PSEN1 |
| CCR5 | MACROH2A1 |
| CAMK2D | PREX1 |
| PELI1 | SLC9A9 |
| ATP2B1 | SCAMP2 |
| ACACA | TOLLIP |
| ATIC | DOK2 |
| KCNMB4 | HLA-A |
| TRIB1 | WDTC1 |
| SH3RF1 | TNFSF13 |
| PTK2 | HEXB |
| GOLGA7B | TNRC18 |
| FLT1 | SLC35E2B |
| PTGIR | LRP10 |
| SRGAP3 | HPS1 |
| ETV5 | VPS39 |
| LPL | ABHD17A |
| KLF4 | TPT1 |
| COX20 | ANXA11 |
| MAP4K4 | CYP27A1 |
| RILPL1 | PLXDC2 |
| PRR5 | IDS |
| OAF | NAGA |
| PDE7A | WBP2 |
| ST6GAL1 | LSP1 |
| IFI44L | IL10RA |
| CALM3 | RPS24 |
| EDNRB | CTSA |
| SLC7A1 | MOB1A |
| ENC1 | UBC |
| DST | TMBIM6 |
| MCM6 | RNASE1 |
| CTTN | MAN2B1 |
| KIF21B | HLA-B |
| ADCK2 | |
| HS3ST2 | |

| | |
|---|---|
| SBF2 | NME4 |
| SCAMP5 | NOL6 |
| SDC2 | NOLC1 |
| SEC31B | NOMO1 |
| SECTM1 | NOMO2 |
| SEL1L3 | NOMO3 |
| SELENOP | NOP10 |
| SENP7 | NOP14 |
| SERINC1 | NOP2 |
| SERPING1 | NR2C2AP |
| SESN1 | NSDHL |
| SESN3 | NSMF |
| SESTD1 | NSUN2 |
| SETDB1 | NUP188 |
| SF3B1 | NUP210 |
| SH3BGRL | NUTF2 |
| SIDT1 | OAF |
| SIDT2 | OGDH |
| SIN3A | ORAI1 |
| SIPA1L1 | ORMDL2 |
| SIPA1L2 | OSGIN1 |
| SIRPB2 | OSM |
| SLC16A10 | OTOA |
| SLC18B1 | P2RY11 |
| SLC22A23 | P2RY2 |
| SLC35E2B | P2RY8 |
| SLC40A1 | PAICS |
| SLC43A2 | PALM2AKAP2 |
| SLC45A4 | PANK3 |
| SLC46A3 | PAPSS1 |
| SLC7A8 | PARVB |
| SLU7 | PC |
| SMAP2 | PCDHGC3 |
| SMIM14 | PCK2 |
| SMOX | PCNA |
| SMPDL3A | PCSK6 |
| SNX29 | PCYT2 |
| SNX6 | PDCD6 |
| SP110 | PDIA6 |
| SPINT2 | PDSS1 |
| SPSB3 | PDXK |
| SPTLC3 | PEA15 |
| SRGAP3 | PES1 |
| SSBP2 | PFAS |
| ST3GAL1 | PFDN6 |
| ST6GAL1 | PFKP |
| ST8SIA4 | PFN1 |
| STAB1 | PGAM1 |
| STK24 | PGK1 |
| STK38L | PGM2 |
| STX16 | PGP |
| SULF2 | PHB1 |
| SYF2 | PHLDA3 |
| SYK | PHLDB1 |
| SYTL3 | PHPT1 |
| TAX1BP1 | PI4K2B |
| TBC1D14 | PIEZO1 |
| TBC1D15 | PIGW |
| TBC1D2B | PIM1 |
| TBC1D3I | PKM |
| TBC1D9 | PKMYT1 |
| TCAF2 | PLEC |
| TENT5A | PLK3 |
| TFPI | PLOD3 |
| TGFBI | PLP2 |
| THBS1 | PLPP3 |
| THEMIS2 | PLXNA1 |
| TLE4 | PMM2 |
| TLR5 | PNP |
| TLR7 | PNPLA6 |
| TLR8 | POLD2 |
| TMEM140 | POLR1A |
| TMEM176A | POLR2L |
| TMEM254 | POLR3H |
| TMEM86A | PPAN |
| TMOD2 | PPARG |
| TNFAIP2 | PPIA |
| TNFRSF21 | PPIB |
| TNFSF10 | PPIF |
| TNFSF12 | PPP1R14B |
| TP53INP1 | PPP5C |
| TPCN1 | PRDX5 |
| TPP1 | PRELID1 |
| TRIM22 | PRMT1 |
| TRIM38 | PRMT5 |
| TSPYL2 | PRPF19 |
| TTC14 | PRR5L |
| TTYH2 | PRXL2A |
| TUG1 | PSEN2 |
| TUT4 | PSMA7 |
| TUT7 | PSMB5 |
| TXLNB | PSMB6 |
| TXNDC16 | PSMC4 |
| TXNIP | PSMD14 |
| UBN2 | PSMD2 |
| ULK1 | PSMG1 |
| UNKL | PTGIR |
| USP30 | PTPN7 |
| UTRN | PTPRE |
| VAMP1 | PTRH2 |
| VCAN | PUF60 |
| VSIR | PUM3 |
| VWA5A | QPRT |
| WASHC4 | RAB10 |
| WBP1L | RAB3IL1 |
| WDFY2 | RAB5IF |
| WDR91 | RAC2 |
| WDTC1 | RAD23B |
| WWC3 | RAN |
| WWP1 | RANBP1 |
| XRN1 | RASGRP3 |
| YPEL1 | RBM28 |
| YPEL2 | RDH11 |
| YPEL3 | RGCC |
| YPEL5 | RGS3 |
| ZEB2 | RHEB |
| ZER1 | RND3 |
| ZFAND1 | RNH1 |
| ZKSCAN8 | RPS19BP1 |
| ZMAT1 | RRP12 |
| ZMYM2 | RRP9 |
| ZNF217 | RRS1 |
| ZNF224 | RSU1 |
| ZNF277 | S100A4 |
| ZNF302 | S100A6 |
| ZNF331 | SAC3D1 |
| ZNF395 | SAE1 |
| ZNF449 | SAPCD2 |
| ZNF532 | SC5D |
| ZNF846 | SCD |
| ZNF862 | SCFD2 |
| ZSCAN18 | SCIN |
| ZSCAN30 | SDC4 |
| | SDF2L1 |
| | SEC11C |
| | SEC13 |
| | SELENOH |
| | SEPTIN11 |
| | SFT2D1 |
| | SFXN1 |
| | SGK1 |
| | SGMS2 |
| | SGPP2 |
| | SH3BGRL3 |
| | SH3RF3 |
| | SH3TC1 |
| | SIGLEC11 |
| | SIGLEC16 |
| | SLC15A4 |
| | SLC16A1 |
| | SLC16A13 |
| | SLC16A3 |
| | SLC17A9 |
| | SLC1A3 |
| | SLC1A4 |
| | SLC1A5 |
| | SLC20A1 |
| | SLC25A1 |
| | SLC25A19 |
| | SLC25A25 |
| | SLC27A4 |
| | SLC2A6 |
| | SLC35B1 |
| | SLC35B4 |
| | SLC39A13 |
| | SLC39A7 |
| | SLC39A8 |
| | SLC43A3 |
| | SLC52A2 |
| | SLC5A3 |
| | SLC6A6 |
| | SLC7A1 |
| | SLC7A5 |
| | SLCO4C1 |
| | SMG5 |
| | SMIM12 |
| | SNCA |
| | SNRNP25 |
| | SNRPD1 |
| | SNRPG |
| | SNTB1 |
| | SNX12 |
| | SOX13 |
| | SPHK1 |
| | SPN |
| | SPR |
| | SPRING1 |
| | SPRY2 |
| | SPTBN1 |
| | SQLE |
| | SREBF1 |
| | SREBF2 |
| | SRM |
| | SRPRB |
| | SRSF2 |
| | SSBP4 |
| | ST14 |
| | ST3GAL6 |
| | STARD4 |
| | STEAP3 |
| | STIP1 |
| | STK40 |
| | STX3 |
| | SUCNR1 |
| | SUN2 |
| | SYNCRIP |
| | SYNE3 |
| | TBC1D2 |
| | TBC1D9B |
| | TBRG4 |
| | TCF19 |
| | TCIRG1 |
| | TEX2 |
| | TFDP1 |
| | TGFBR1 |
| | TGM2 |

RRM2
PGA4
TK1
IL6
SIGLEC11
GSDMA
KIF20A
FLT1
TYMS
KCNJ15
MCM10
MKI67
DDIAS
CCL3L1
BIRC5
ANKRD22
LOC102724951
LOC102724219
LOC102724843
CCL13
SERPINE1
TNC
MYBL2
CD1B
TMEM45B
TACSTD2
FAM111B
TNFSF12-TNFSF13
E2F8
LGALS2
DTL
PGA3
CLEC5A
PGA5
MCEMP1
TREM1
RPS10-NUDT3
CXCL5
PPBP
CCL7

ZMAT1
C2orf42
DCP1A
SIRPB1
DNM2
KLHL21
MYO9B
MIR138-2
GYPC
JOSD1
RC3H2
SUPT4H1
TTC39C-AS1
CBR3-AS1
FNDC3B
H3F3B
RUNX1
ATP6V0C
ADPGK
COPA
EIF1
LMO4
NUP98
PNRC1
ZNF221
NDEL1
TPRA1
CDC42SE1
FAM168B
LYRM1
NFKBIA
PIK3AP1
SLK
WASF1
NDST2
CD163L1
ESYT2
HIST1H2AH
IVNS1ABP
LONP1
PLTP
GARS
LAMTOR4
LINC00963
KLF10
SH3BP2
SRSF3
FAM114A2
NCF2
USP47
ARMC5
ATXN1L
CCDC114
KIAA0895L
NR1D2
PHC3
LOC151475
PSME4
RANBP10
RPS12
ZSCAN18
SLC7A5P2
DEFB1
PJA2
KLF4
STX18
EIF5
FLJ32255
TTC9C
SLF2
TOB1
TP53INP2
RBM15B
RNF19B
CEBPB
GRK6
MALAT1
MIDN
NPL
SHB
WSB1
ZBTB1
AGAP3
ITGB1
LINC01612
IER5L
NDUFB6
PIK3R3
SNHG19
SRSF6
TRA2A
CTSZ
GIMAP4
HEATR5A
HSDL2
IDI1
MYCBP2
SERPINB1
SSU72
ATF7IP
EAPP
HGSNAT
KIDINS220
LONRF3
RBMX
TMX1
TSPYL2
UBE2A
UFD1L
AP3S1
BLOC1S6
SEMA7A
ATG4B
AVL9
COMMD5
OIP5
RNU5A-1
SUMO3
LARP7
NMT1
RBM23
RUFY1
RAB7B
TARS
ATP1B3
ATP6V0B
CSRNP1
PAXIP1
TCF4
PABPN1
RAP2B
TMSB10
CTSL
HEATR5B
HEIH
EIF4A1
ERF
MFSD1
NADK2
NINJ1
PPIA
SWT1
UNC50
AMD1
MRPS30
CHFR
TMBIM6
SLC16A3
MAN2B2
SLC38A2
FAM110A
KMT5A
SLBP
SMC5
FGR
HIF1A
MRAS
CDKL3
EXOC7
HIST1H2AE
LOC440700
PRMT9
SERPINB6
TAF8
TNFAIP6
FMO4
PXDC1
MAPK8
NUP37
ARPC2
HPS3
S100A12
TPCN1
ANKRD13A
CMC2
FAS
LINC01136
MGAT1
NSMCE2
OSER1-AS1
PHACTR1
PLRG1
RPL3
SLC25A33
SLC35A5
SRSF2
TYMP
ARSD
RAB5C
ARHGAP26
CCDC47
DNAAF5
TOB2
CSTB
SLC20A1
SF1
RNU5E-1
MIR6125
SHMT2
TMEM14C
LRCH4
LINC01134
GSTO1
HNRNPAB
ODF2
RPL12
MICALL2
EIF3B
DLGAP1-AS1
MED18
TBC1D15
MKRN2
HMGB2
ATF6
DNAJB1
ELL2

NABP1
LOC102723728
KANK2
IVNS1ABP
MYO1E
SLC45A3
BNC2
PPARG
CALCRL
FZD1
SESN3
SPDL1
OTOA
AK4
PAICS
GADD45B
NREP
UCP2
DENND11
C1orf112
P2RY1
EPB41L2
RASGEF1B
GPX3
HDAC9
CCRL2
GPR161
LILRA5
NUSAP1
FMNL2
IL4I1
DDAH2
CCL8
STARD4
FKBP5
LGMN
TRIM47
APOC1
RASGRF1
ECE1
AQP9
IL18
LTBP2
DHCR24
KIF20B
IL1R2
CTDSPL
DDR1
NIBAN1
GYPC
SPP1
TMEM106C
TCF7L2
IFIT2
GINS1
RHOB
SLAMF6
ITIH4
CD28
SLC1A3
PDE2A
SLC18B1
CENPN
TNC
TGFA
HPGDS
NLN
OBSCN
C1orf21
FHIT
DMD
PLXNA1
SLCO4A1
FAR2
MT2A
GPBAR1
NOL4L
IL1RN
RASGRP2
CDCA7L
RTN4R
MYO10
FABP5
SQLE
RFC3
DHRS3
STMN1
SPTBN1
RAI14
LDLRAD3
SELL
NEDD9
STRIP2
DCSTAMP
ARHGAP32
EPS8
RGS16
GIMAP5
CDKN2B
BUB1
PTTG1
HPSE
FADS2
CD209
RUBCNL
PRR5L
LRRC4
LILRB5
RGS3
AKAP6
SYT1
NR1D1
FSCN1
CKS2
SPTLC3
RAB3IL1
MYO1B
SLC40A1
ITGA6
POU2F2
KLF10
METTL7B
SLC28A3
SEL1L3
KITLG
MAPK12
HSF4
NMRK2
IGFBP4
P2RX7
MYC
FAM174B
SLC35F2
PMAIP1
GNG2
STEAP3
CENPU
MAD2L1
PLPP3
MRC2
FOXM1
SLC22A23
KLHL13
PALM2AKAP2
TLE1
ADRB2
SDS
HRH1
SHCBP1
NRGN
HIVEP3
JPH4
DNASE2B
CARD11
BCAR3
SDC3
KIF2C
CCL3
GAL3ST4
SCN9A
CCNB1
FRMD3
MELK
FN1
NAV1
ALS2CL
TMEM273
GINS2
HGF
MTHFD1L
PRLR
CDCA5
CD22
SYT17
CHST2
TLCD3A
CCNB2
EYA2
CDCA2
PCED1B
KCNJ5
CDC20
CCL4L1
ABCA6
INKA1
CCNA2
GPRC5B
CCL4
NAV2
GTSE1
AURKB
SLC37A3
KIAA1549
DHRS9
ANLN
DDIAS
CD200R1
EGR3
TNFSF15
TPX2
AMIGO2
MRO
KIF4A
SAPCD2
CEP55
HMMR
MCC
TROAP
TOP2A
CCL3L1
CDT1
KIF23
UHRF1
KIFC1
ITGAD
NIPAL4

TIMM13
TIMM17A
TIMM23
TLNRD1
TM4SF19
TMC8
TMED3
TMED9
TMEM250
TMEM97
TMIGD3
TMX2
TNF
TNFRSF12A
TNFSF15
TNS1
TOLLIP
TOMM40
TOMM40L
TOMM5
TP53I11
TPI1
TRAPPC2L
TREM2
TRIB3
TRIM47
TRIM65
TRMT61A
TRPV2
TRUB2
TSPO
TTLL12
TUBA1B
TUBB
TUBB2A
TUBB4B
TUBB6
TUBG1
TWF2
TXNDC5
TXNRD2
UAP1L1
UBASH3B
UBE2S
UBL4A
UCK2
UHRF1
UNG
UQCC2
UQCRC1
URB1
UROD
USP5
UTP4
VAC14
VARS1
VCL
VCP
VDAC1
VDR
VOPP1
VPS18
WDR1
WDR12
WDR4
WDR46
WDR77
YWHAG
YWHAH
ZDHHC16
ZNF703
ZNF787

| | |
|---|---|
| *IDH1* | *NR4A1* |
| *INPP5D* | *WIPF3* |
| *LOC101929210* | *ABCA9* |
| *PPARGC1A* | *E2F1* |
| *SGTA* | *FBLIM1* |
| *SOD2* | *TK1* |
| *STAT3* | *IRAG1* |
| *TMEM123* | *PLK1* |
| *NASP* | *ZNF367* |
| *SGSM2* | *NCAM1* |
| *USO1* | *CCL24* |
| *STRIP1* | *FBLN5* |
| *CCNYL1* | *CCL4L2* |
| *CTSB* | *PCDH12* |
| *GOSR2* | *CLSPN* |
| *RGS1* | *CDK1* |
| *MESDC1* | *TNFSF18* |
| *CHCHD7* | *HJURP* |
| *EIF2S2* | *PRSS23* |
| *NAMPT* | *MKI67* |
| *RBM38* | *BOLA2B* |
| *NSDHL* | *PLXDC1* |
| *ZDHHC4* | *DTL* |
| *UBXN4* | *CX3CR1* |
| *KPNA4* | *NEK2* |
| *PIM3* | *HELLS* |
| *HOOK3* | *BUB1B* |
| *MIR4530* | *FCER2* |
| *ARHGDIA* | *CDCA7* |
| *GPR84* | *BIRC5* |
| *INSIG1* | *NDC80* |
| *SETD4* | *NCAPG* |
| *PNKD* | *KIF14* |
| *ARRDC3* | *UBE2C* |
| *ACTB* | *P3H2* |
| *ARSB* | *FFAR2* |
| *CRYL1* | *SPRY1* |
| *FBXL14* | *TRIP13* |
| *GNAI3* | *SYT6* |
| *ID3* | *OLFML3* |
| *PIGP* | *SKA3* |
| *MIR7845* | *CKAP2L* |
| *CYTH2* | *KIF20A* |
| *NEUROD4* | *DEPDC1* |
| *YLPM1* | *PHGDH* |
| *MKLN1-AS* | *PCLAF* |
| *EIF3D* | *APOBEC3B* |
| *LINC01578* | *MYBL2* |
| *SEC1P* | *FAM111B* |
| *SLC7A5* | *HAMP* |
| *UCHL3* | *MAMDC2* |
| *C11orf71* | *CCNE2* |
| *CNP* | *CSGALNACT1* |
| *HMGB1* | *DIAPH3* |
| *INTS6* | *BIRC7* |
| *LOC100996437* | *PKMYT1* |
| *NFKBID* | *EXO1* |
| *SARAF* | *DLGAP5* |
| *TTN-AS1* | *CDC6* |
| *FAM210A* | *KIF18B* |
| *ALK* | *APOBEC3A* |
| *PTPN1* | *SPC25* |
| *PTDSS1* | *AFF3* |
| *ARL6IP6* | *MCM10* |
| *GNL3* | *TYMS* |
| *MIR4444-1* | *RRM2* |
| *ARID5A* | *CDC45* |
| *CD164* | *E2F8* |
| *GAPDH* | |
| *SUSD6* | |
| *NFAM1* | |
| *TMEM185B* | |
| *FAM219A* | |
| *CHN2* | |
| *DNAJC3* | |
| *HDAC4* | |
| *LMNA* | |
| *SUPT5H* | |
| *ZNF706* | |
| *LIAS* | |
| *HIST2H2AC* | |
| *IFRD1* | |
| *ANKHD1-EIF4EBP3* | |
| *IFI30* | |
| *C19orf53* | |
| *SLFN5* | |
| *TMEM187* | |
| *CDKN2AIP* | |
| *PRCP* | |
| *SAT1* | |
| *SLC7A7* | |
| *SLC8B1* | |
| *SRRM2-AS1* | |
| *TBC1D22A* | |
| *AKIRIN2* | |
| *RPL10A* | |
| *HPCAL1* | |
| *TUBB4B* | |
| *ZNF592* | |
| *IRF8* | |
| *IBA57* | |
| *PURB* | |
| *XBP1* | |
| *LINC01353* | |
| *NCR3LG1* | |
| *SGSH* | |
| *NSUN6* | |
| *LOC101926933* | |
| *CSTF1* | |
| *ZNF628* | |
| *RRM2B* | |
| *FBXO28* | |
| *RRNAD1* | |
| *RNVU1-18* | |
| *NOTCH1* | |
| *LOC101927780* | |
| *MOCS3* | |
| *MCTP1* | |
| *LOC100289230* | |
| *SSR1* | |
| *SSH1* | |
| *RNVU1-19* | |
| *TRA2B* | |
| *SIGLECL1* | |
| *NRIP1* | |
| *AHR* | |
| *NBR2* | |
| *SFT2D1* | |
| *WBP1L* | |
| *GNB1* | |
| *KANSL3* | |
| *RILPL2* | |
| *LINC01094* | |
| *GABARAPL1* | |
| *MPHOSPH9* | |
| *RABGAP1* | |
| *ARHGAP26-IT1* | |
| *RNU4ATAC* | |
| *NPAT* | |
| *SDCCAG3* | |
| *ZNF767P* | |
| *HBP1* | |
| *MGC16275* | |
| *SNORD3A* | |
| *FBXO31* | |
| *SKOR1* | |
| *HECTD1* | |
| *CTBP1* | |
| *CACYBP* | |
| *DENND4A* | |
| *ZNF143* | |
| *TM2D3* | |
| *PHLDA1* | |
